# Lipidomics Reveals Cell Specific Changes During Pluripotent Differentiation to Neural and Mesodermal Lineages

**DOI:** 10.1101/2024.12.31.630916

**Authors:** Melanie T. Odenkirk, Haley C. Jostes, Kevin R. Francis, Erin S. Baker

## Abstract

Due to their self-renewal and differentiation capabilities, pluripotent stem cells hold immense potential for advancing our understanding of human disease and developing cell-based or pharmacological interventions. Realizing this potential, however, requires a thorough understanding of the basal cellular mechanisms which occur during differentiation. Lipids are critical molecules that define the morphological, biochemical, and functional role of cells. This, combined with emerging evidence linking lipids to neurodegeneration, cardiovascular health, and other diseases, makes lipids a critical class of analytes to assess normal and abnormal cellular processes. While previous work has examined the lipid composition of stem cells, uncertainties remain about which changes are conserved and which are unique across distinct cell types. In this study, we investigated lipid alterations of induced pluripotent stem cells (iPSCs) at critical stages of differentiation toward neural or mesodermal fates. Lipdiomic analyses of distinct differentiation stages were completed using a platform coupling liquid chromatography, ion mobility spectrometry, and mass spectrometry (LC-IMS-MS) separations. Results illustrated a shared triacylglyceride and free fatty acid accumulation in early iPSCs that were utilized at different stages of differentiation. Unique fluctuations through differentiation were also observed for certain phospholipid classes, sphingomyelins, and ceramides. These insights into lipid fluctuations across iPSC differentiation enhance our fundamental understanding of lipid metabolism within pluripotent stem cells and during differentiation, while also paving the way for a more precise and effective application of pluripotent stem cells in human disease interventions.

## Introduction

Animal and cellular disease models have a history of advancing insights into complex biological processes and accelerating the discovery of therapeutic interventions. Specifically, animal and cell lines have significantly advanced disease mitigation efforts for polio, various influenza strains, cancer subtypes, Ebola, and COVID-19.^1–5^ However, translational challenges from animal models to humans have limited effectiveness^.6, 7^ To date, approximately 90% of clinical trials fail to produce effective treatments for humans. It is therefore evident that existing disease models, despite their contributions, are not always sufficient for accurately predicting human responses.^6–8^ Pluripotent stem cells (PSCs) have recently bridged this challenge by offering a heterogenous cellular environment that more accurately reflects complex tissues. Specifically, both embryonic stem cells (ESCs) and induced pluripotent stem cells (iPSCs), exhibit self-renewal and differentiation, two key abilities allowing these cells to develop into defined cell types. Pluripotent stem cells have already demonstrated a revolutionary potential to overcome existing translational challenges of previous disease models.^9, 10^

The differentiation of PSCs to mature cell types results in drastic morphological, biochemical, signaling, and functional changes, which need to be understood to optimize their use as effective disease models.^11^ Lipids, a class of nonpolar small molecules, have shown critical, cell-specific composition and function.^11, 12^ PSCs exhibit an abundance of highly unsaturated lipids, a phenomenon potentially related to the plasticity of these cells to generate a myriad of specialized, mature cell types.^13^ Undifferentiated PSCs, similar to cancerous cell types, rely on activated glycolysis and *de novo* fatty acid synthesis as their primary energy source.^11, 14^ Moreover, studies evaluating PSC development into mature cells types have demonstrated massive lipidome alterations for 5 out of 8 lipid categories including the glycerophospholipids, glycerolipids, sphingolipids, sterols, and fatty acyls.^11, 14–16^ Taken together, preliminary metabolic work on PSCs has identified lipids as vital molecules for determining cell fate and survival, making the understanding of their differentiation a critical prerequisite knowledge for PSC disease model success.^16^

To date, studies investigating lipidome alterations associated with stem cell differentiation have focused on a singular cell fate, however the question of whether previously observed lipidome changes reflect general PSC differentiation mechanisms or are instead cell-type specific has remained unanswered.^17^ A comprehensive and longitudinal evaluation of lipidome changes during PSC commitment to specific cellular lineages is therefore essential to uncover conserved and unique lipid alterations associated with cellular differentiation. Herein, we longitudinally monitor lipidomic alterations during the differentiation of neural and mesodermal lineages from iPSCs using established differentiation assays. Lipidome changes were tracked as cells transitioned from pluripotency to intermediate lineages and then to defined cell types, in order to characterize conserved and unique lipid biomarkers of iPSC differentiation.

## Results and Discussion

To evaluate how lipid species changed during iPSC differentiation to neural and mesodermal lineages, lipidomic assessments were performed on differentiated derivatives originating from the NCRM-5 human iPSC line.^18^ Neural cell differentiation occurred over 42 days using a modified rosette-based differentiation and isolation method.^18, 19^ Mesodermal cells were differentiated over 21 days following an established differentiation protocol^.20^ For both lineage-directed differentiation methods, samples were collected at specific developmental timepoints to define molecular differences between cell states and lineages (**Figure 1**). Lipids were isolated from cells through a modified Folch extraction before being analyzed with the multidimensional LC-IMS-CID-MS platform^.21, 22^ A total of 350 unique lipid identifications were observed during the neural differentiation process, while 453 unique lipid identifications were detected during mesodermal specification. Initially, we explored the lipidome alterations longitudinally across each cell type and then compared the individual lipid profiles for the neural and mesodermal fates to identify unique and conserved lipidomic alterations.

**Figure 1.**
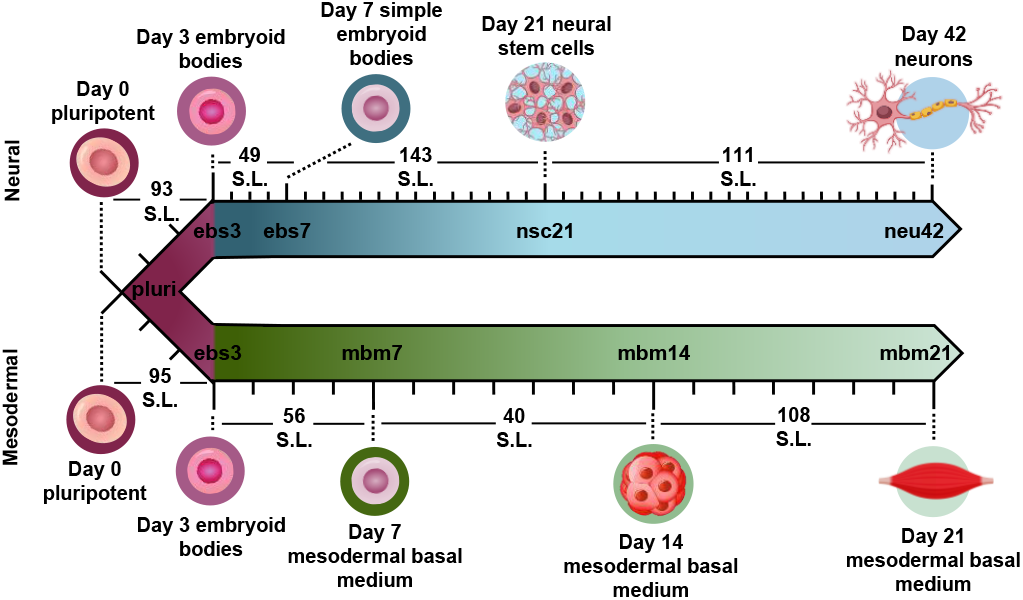
Overview of stem cell differentiation timepoints and the number of statistically significant lipids (S.L.) observed for each timepoint comparison. Stem cells were cultivated into neural (top) and general mesodermal (bottom) cells from an iPSC stem cell line derived from CD34+ cells from cord blood.^18^

### Neural Lineage Commitment

For neural differentiation, iPSCs were differentiated over a period of 42 days, with cellular harvesting at 4 significant timepoints during development: pluripotent cells (day 0; pluri), embryoid bodies (day 3; ebs3 and day 7; ebs7), neural stem cells (day 21; nsc21), and neurons (day 42; neu42).^18, 19^ Successive timepoint comparisons illustrated that of the 350 unique lipids detected, 207 were statistically significant (α=0.05) in at least one timepoint (with many statistically significant in multiple timepoints). Specifically, in day 3 embryoid bodies (ebs3) compared to pluripotency (pluri), 93 lipids were statistically altered (**Figure 1**). These differences, however, decreased in day 7 (ebs7) vs. ebs3, where only 49 lipids were statistically different. However, following neural stem cell formation at day 21, an influx of lipidome changes were observed with 143 lipid species being significantly changed relative to the ebs7 timepoint. The final comparison of neu42 vs. nsc21 yielded a total of 111 significant lipid fluctuations. Interestingly, a similar number of lipids exhibited increased and decreased expression across all timepoint comparisons, with a slight bias towards increased expression (49 upregulated, 44 downregulated) in the comparison of neu42 vs. nsc21. Additionally, most altered lipids for each successive timepoint comparison were also significant in at least one other comparison. Taken together, these results demonstrate that the lipids were significantly altered during cell differentiation at all measured timepoints.

To more thoroughly understand the lipid changes occurring during differentiation, lipid class and head group trends were assessed using the Structural-based Connectivity and Omic Phenotype Evaluation (SCOPE) cheminformatic toolbox.^23^ SCOPE utilizes lipid structural information to link species to their biological relationships which can then be visualized using hierarchical clustering and grouped heatmaps. A circular dendrogram incorporating hierarchical clustering information is subsequently used to evaluate significant lipid changes within the lipid classes for each timepoint comparison (**Figure 2A**). This dendrogram was created by plotting the observed log_2_FC of all detected lipids using a red/blue gradient to depict the magnitude and direction of change among the significant lipid species. Several trends were observed in this study, suggesting the involvement of multiple lipid classes in iPSC maturation into neurons^.23^ Triacylglycerols (TGs), for example, showed differential expression patterns based on lineage timepoints as did several other lipid classes. For example, in ebs3, TGs increased relative to pluripotent cells, a finding that may reflect the synthesis of TGs for use as a cellular energy source.^24^ Notably, free fatty acid (FFA) upregulation was also observed at this timepoint. This trend was reversed and both TGs and FFAs were mainly decreased following cell differentiation from day 21 to day 42, possibly due to their use in different energetic processes.^11^

**Figure 2.**
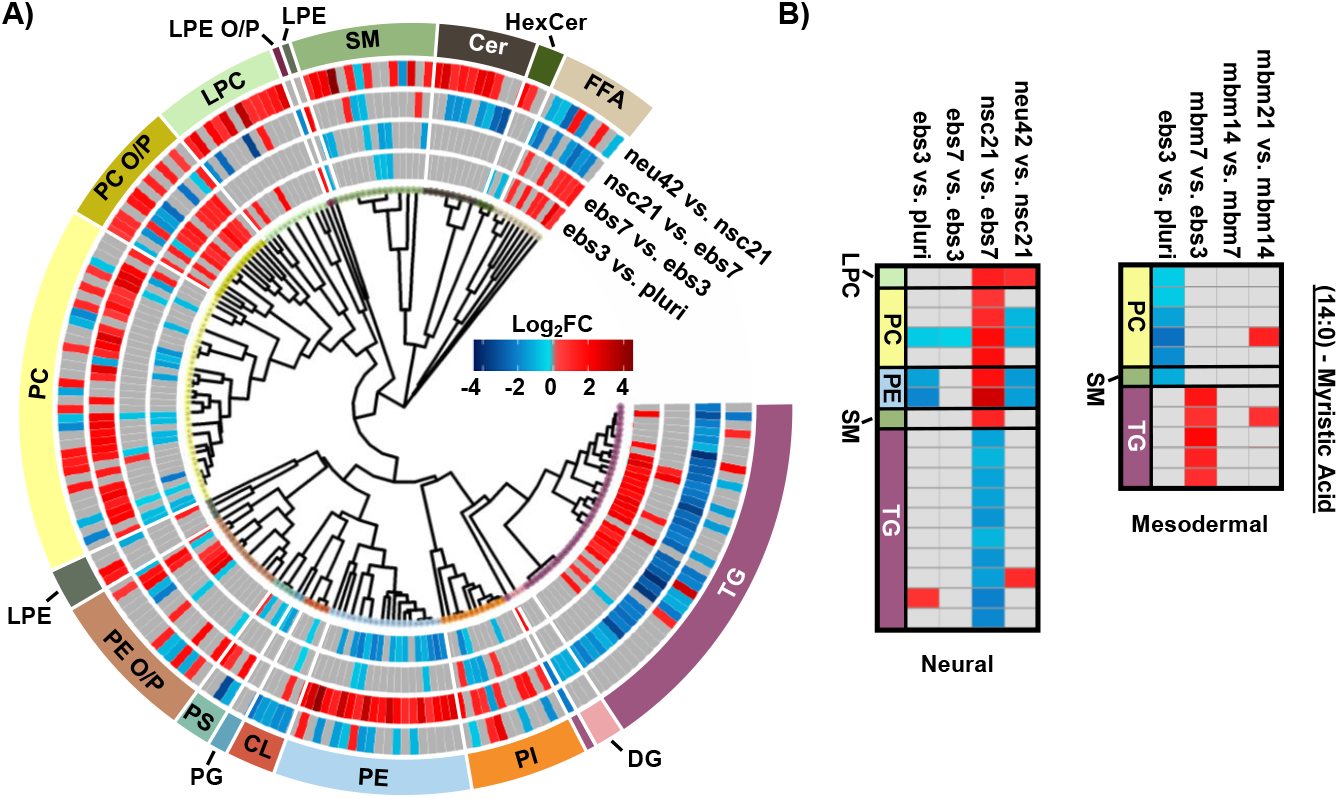
Neural stem cell line lipid alterations. A) Circular dendrogram of significant lipid changes overtime. For each timepoint, 4 biological replicates were used (n=4), except neu42 where n=5. B) Heatmap of significant fatty acyl changes for myristic acid (14:0) in the neural (left) and mesodermal (right) cell lines. Red and blue denote the log2 fold change of significant species that are up- or downregulated. Grey denotes lipids with no statistical significance.

Evaluation of other lipid species changing during differentiation also highlighted additional trends. For example, phosphatidylethanolamine (PE), phosphatidylcholine (PC), and phosphatidylinositol (PI) species had an opposite trend to TG and the FFA classes. Specifically, lower abundances for these select phospholipids were observed in early differentiation of iPSCs and increased upon neural stem cell specification (nsc21). Phospholipids are integral species in lipid bilayers and cellular signaling processes that render them of great importance in neural cell proliferation.^16^ However, comparison of neu42 and nsc21 timepoints demonstrate both increased and decreased expression of species within these phospholipid classes that may suggest a preferential expression of specific phospholipid species in neurons compared to neural stem cells. Additionally, alkyl and alkenyl phosphatidylcholine (PC O/P), alkyl and alkenyl phosphatidylethanolamine (PE O/P), and lysophosphatidylcholine (LPC) lipids exhibited increased abundance over time except in the nsc21 vs. ebs7 timepoint comparison. Alkyl (O) and alkenyl (P) ether phospholipids are a minor composition of the human glycerophospho-lipidome that have an enhanced capability to release their *sn*-2 fatty acyl for lipid mediator synthesis following hydrolysis.^25, 26^ Neural lipidome studies have linked decreases in plasmalogens with oxidative stress that is associated with neurodegenerative diseases such as Alzheimer’s disease.^17^ A decrease in cardiolipins (CL) was also observed within the neu42 vs. nsc21 timepoint comparison, potentially related to mitochondrial function reflective of oxidative processes as was observed with the plasmalogen upregulation.^27^ Additionally, upregulation of sphingomyelin (SM), ceramide (Cer) and hexose ceramide (HexCer) lipids was observed within differentiated neurons. A previous study comparing iPSCs to iPSC-derived neurons also observed an increase in Cers and attributed this to their role in neuronal differentiation.^28, 29^ Sphingolipids also have a documented influence on driving pluripotent stem cells to a neural fate and increasing membrane fluidity via the SIRT1 enzyme, whereby sphingolipid deficiency has been linked to impaired neural differentiation.^30^ Moreover, glucose/galactose glycosylated sphingolipids have a significant influence on axonal growth, potentially explaining their influx in the neu42 vs. nsc21 timepoint comparisons.^31^

In addition to the head group analyses, fatty acid (FA) trends were examined by plotting composition changes for each FA group. During neural differentiation, 18 lipids containing a 14:0 moiety, commonly known as myristic acid (MA), were found to be statistically significant (**Figure 2B**). MA is broadly incorporated into phospholipids to contribute to cell membrane structure.^32^ Additionally, MA is involved in myristoylation which is a co- and post-translational lipidation modification in which the acyl group is covalently attached to the amino terminus (N-terminus) of cellular proteins by N-myristoyltransferase (NMT)^.32, 33^ Myristoylation enables a diverse range of biological functions such as protein binding in membranes, protein-protein interactions, signal transduction, cellular transformation, and subcellular localization. Within the nervous system, myristoylation of synaptic proteins is important for synaptic plasticity.^34^ Furthermore, MA has been previously demonstrated to promote neural stem cell proliferation and differentiation *in vitro* and *in vivo*, generating great interest in further studies regarding its use in the treatment of neurological disorders^.35-38^ The preferential storage of 14:0 FAs as phospholipids at the nsc21 vs. ebs7 timepoint comparison could be an indication of enzymatic phospholipid remodeling, as this is one source of MA within cells^.34^ The subsequent downregulation of 14:0 containing phospholipids and FFAs at the final neu42 vs. nsc21 timepoint comparison may suggest the use of MA in neural stem cell differentiation. The same dysregulation trends were observed for both TG and phospholipid species containing a 16:0 and 16:1 moiety, which are more commonly known as palmitic acid (PA) and palmitoleic acid. PA is an energy source and similar to MA, serves in protein lipid modifications through a reversible palmitoylation process^.39, 40^ Protein S-palmitoylation is the most common acylation in eukaryotic cells and aids in the regulation of neuronal protein trafficking, signaling, and function. Specific

### Mesodermal

palmitoylation targets have also been shown to help regulate synaptic plasticity.^41^ Palmitoylation aids in the regulation of neuronal protein trafficking and function. Taken together, these trends indicate that short chain FA containing lipid species may contribute to the development of neurons. Additional follow-up studies comparing protein post-translational myristoylation and palmitoylation at these stages of differentiation could further demonstrate these observations.

### Mesodermal Stem Cell Lineage

To encapsulate the uniqueness of lipid changes in neural cell development from iPSCs relative to general iPSC differentiation, a mesodermal differentiation protocol was also performed on NCRM-5 iPSCs. As mesodermal stem cell differentiation occurs more rapidly than neuron development (21 days vs. 42 days for neurons), mesodermal lineage differentiation timepoints were collected synonymous to the initial, intermediate, and terminal cell differentiation steps in the neural analyses. Samples collected included pluripotent cells (day 0; pluri), embryoid bodies (day 3; ebs3), mesodermal basal medium (day 7; mbm7 and day 14; mbm14) and general mesodermal differentiated cells (day 21; mbm21). Overall, 453 unique lipids were detected in the mesodermal lineage which was 103 more than the neural study (350 unique lipids). Interestingly, 191 of the 453 lipids were statistically significant in at least one timepoint, which is less than the neural study (207 statistically significant from 350 detected).

Investigation into the successive timepoint lipid changes showed a total of 95 lipids were significantly altered when comparing the ebs3 vs. pluri timepoints (**Figure 1**). Intermediate timepoint comparisons, however, showed a decrease in the number of significant lipidome alterations, with only 56 lipids changing between the mbm7 vs. ebs3 timepoints and 40 lipids between the mbm14 vs. mbm7 timepoints. The greatest lipid fluctuations were observed between mbm21 vs. mbm14 where 108 species were statistically significant. Similar to the neural lineage data, fluctuations were most common at the initial and final timepoints analyzed. Uniquely, however, the mesodermal cell line included slightly more lipid downregulation when compared to the neural differentiation assays. Meanwhile, in the mbm7 vs. ebs3 comparison, 82% of significant lipids were upregulated. From this data, we also observed an increase in the number of uniquely varying lipid species at later timepoint comparisons, similar to the neural analyses.

To explore the mesodermal lineage in greater detail, SCOPE^23^ was again used to navigate both head group and FA trends in significant lipidome alterations of the mesodermal lineage. First, class-based lipid trends were explored with hierarchical clustering (**Figure 3A**). TGs and FFAs showed similar expression patterns and directionality to the neural lineage, with initial upregulation and then decreased expression at the 21-day timepoint. In contrast to neural differentiation, we observed influx and subsequent loss of PC O/P alkyl ether and plasmalogen species during mesodermal lineage formation while increased expression was observed in both the first and final stages of the neural lineage differentiation timepoints. This trend, however, was not as apparent for PE O/P species as both up- and downregulation were observed within both early and late differentiation comparisons. Additionally, the trend of PC O/P-lipids changes over time was opposite of other phospholipid (PI and PE) classes. The fewer significant lipids for each timepoint in the mesodermal data, however, caused the head group trends to be limited relative to the hundreds of lipid fluctuations in neural stem cell formation. While different lipidomic fluctuations may arise for defined mesodermal-derived cell types, the purpose of these experiments was to provide a germ-layer comparison for the neural lineage.

**Figure 3.**
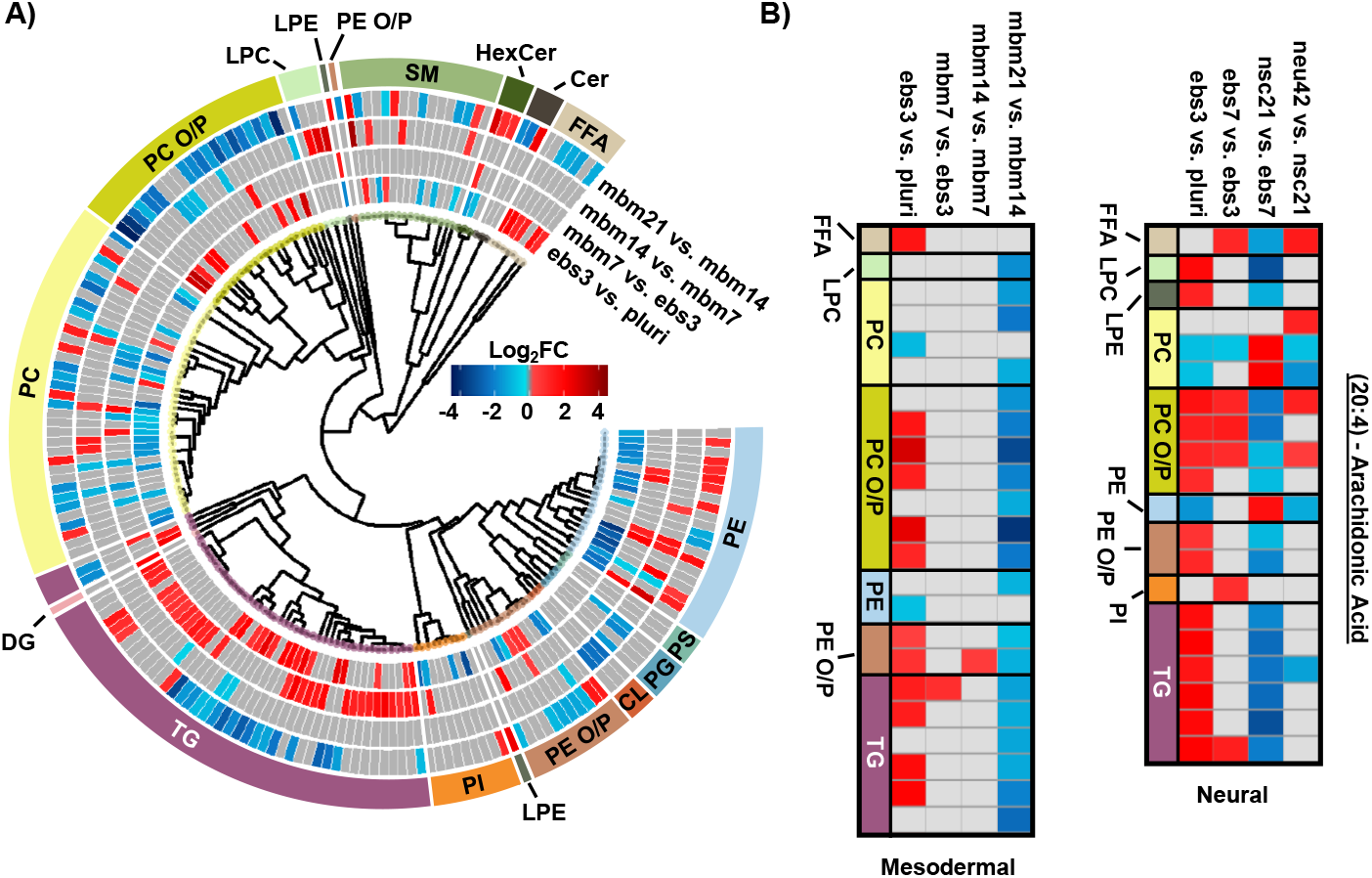
Overview of statistically significant lipid alterations during mesodermal stem cell differentiation. A) Circular dendrogram of the log2FC values observed for the statistically significant lipid species in subsequent differentiation steps with n=4 for each timepoint. B) Fatty acyl heatmap of the log2FC values of arachidonic acid (20:4) in the mesodermal (left) and neural (right) cell lines.

The FA composition changes were next evaluated for the mesodermal lineage. We observed a time specific trend for 23 lipids with a 20:4 moiety (**Figure 3B**). The FFA 20:4, also known as arachidonic acid (AA), has been shown to hyperpolarize cells, contribute to bone metabolism, and serve as a precursor for a series of eicosanoid lipid mediators that facilitate cellular signaling and inflammatory responses.^42–44^ The early accumulation of AA observed here has previously been demonstrated by the preferential accumulation of AA-CoA in cell types of the mesodermal lineage^.42^ The downregulation of AA lipids observed in the final mbm21 vs. mbm14 timepoint comparison was observed in both glycerophospholipid and glycerolipid species. Glycerophospholipids have been linked as the primary source of AA through the cleavage of this fatty acyl moiety with phospholipase enzyme 2 (PLA2) through a hydrolysis reaction.^45^ AA can then be used to synthesize eicosanoid lipid mediators which facilitate cell proliferation and stem cell differentiation.^46^ The downregulation of glycerolipids containing AA has been suggested through a more novel mechanism of lipid droplet-associated TGs that requires the presence of an ATGL enzyme.^47, 48^ Notably, ATGL is highly expressed within white and brown adipose tissue, mature cell types that originate from the mesodermal layer, that may support the feasibility of this mechanism in addition to previous findings of lipid droplet formation during iPSC development.^24, 48^ The loss of AA-containing TGs is especially interesting relative to neutral lipid storage diseases where an accumulation of TGs has been well-defined across many tissue types.^48^ While complex lipid species containing 20:4 moieties were downregulated, no supporting accumulation of 20:4 as a FFA was observed. However, this could reflect the downstream synthesis of AA-metabolites.

When assessing 20:4 FA-containing lipids trends in the neural lineage, a similar trend was observed to the mesodermal, with an initial upregulation of lipids containing 20:4 and downregulation at later time points. However, the time point of differentiation was unique (**Figure 3B**). Comparison of the 14:0, 16:0, and 16:1 carbon FAs from the neural study to the mesodermal cell line illustrated largely unique trends (**Figure 2B**). Specifically, the phospholipid species were initially downregulated in both, though to a much greater extent within the mesodermal differentiation. Subsequent upregulation in the nsc21 vs. ebs7 time point was unique to the neural lineage. Furthermore, TG lipids containing 14:0, 16:0, or 16:1 FAs were largely upregulated in the comparison of mesodermal ebs7 vs. ebs3. Alternatively, neural assays showed a dramatic loss of TGs at the nsc21 vs ebs7 timepoint. This suggests that the mechanisms utilizing short chain FA containing lipids in neural cells are unique, particularly for TG species.

### Lipidomics Results Across Cell Lines

To understand the unique lipidome changes within each cell type, direct comparisons of the differentiation timepoints were performed. Here, hierarchical clustering was used to examine class-based lipid trends within each lineage for the ebs3 and final differentiation timepoints compared to the pluripotent stage (**Figure 4**). For the ebs3 vs. pluri timepoint comparison, we observed similar general class trends for each cell type; however, we did notice unique lipidome compositions. This is most notable in the number and identity of the lipid species that are either up- or downregulated within each class. As this timepoint reflects the transition of pluripotent cells to embryoid bodies, it is hypothesized that the differences in lipidome composition are triggered by the altered biological and physical growth conditions used to induce loss of pluripotency and induction of embryoid body formation. When examining the final stages of differentiation compared to the initial pluripotent stage, there is clear evidence of distinct lipidome changes within each lineage directed method. While there are some shared class-based shifts, such as the downregulation of CLs, significantly changed lipid species typically exhibit opposite trends within the two differentiation protocols. For example, PCs demonstrated some of the greatest abundance shifts when comparing each endpoint to the pluri stage (**Figure 4**). In the neural lineage, many of the identified PCs increased in abundance in the neu42 timepoint. This trend was also observed in a previous study which sought to characterize lipid composition differences between human iPSCs and iPSC-derived neurons.^28^ The noted abundance change may be due to the role of phospholipids in cellular signaling processes, which makes them particularly important in neural cell proliferation.^16^ Interestingly, this trend is not present in the mesodermal lineage. Rather, a general decrease in abundance of PCs is observed between the end stages. This is also consistent with a previous study evaluating broad class-based differences between stem cells and mouse embryonic fibroblast cells, a potential fate for mesodermal stem cells.^49^ While the functional implication of the decreased abundance in the mesodermal lineage remains unclear, these results emphasize the importance of lipid composition in cell function and fate, and the uniqueness of these changes in different cellular lineages.

**Figure 4.**
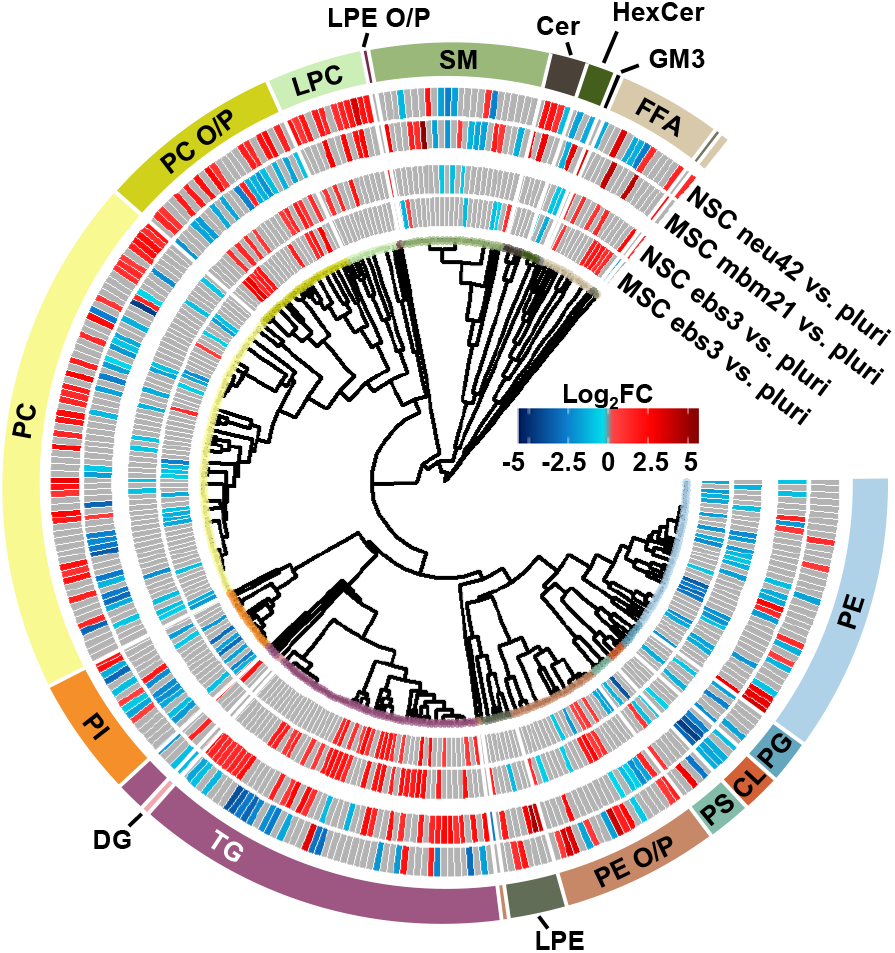
Mesodermal stem cell differentiation relative to the neural lineage. Circular dendrogram of the log2FC values observed for significant lipid species in the first and final differentiation timepoints of each lineage. For each timepoint n=4, except NSC neu42 where n=5.

### Experimental

#### Human iPSC Culture and Differentiation

The NCRM-5 human iPSC line (derived from cord blood CD34+ cells from a male donor) was a kind gift from the iPSC Core Facility, National Heart, Lung, and Blood Institute, located in Bethesda, MD.^18^ iPSCs were maintained in StemMACS iPS-Brew XF medium (Miltenyi Biotec) on Matrigel hESC-qualified matrix (Corning). Neural induction utilized a modified rosette-based method as previously published.^18, 19^ For neural induction, embryoid bodies were generated in Aggrewell 800 plates (Stem Cell Technologies) following manufacturer recommendations in the following media: DMEM, B-27 supplement with vitamin A, N2 supplement, LDN193189 (100 nM), SB431542 (10 µM), FGF2 (10 µg/mL), EGF (10 µg/mL), glutamine (2 mM), and penicillin-streptomycin (50 µg/mL). Y27632 (10 µM; Reagents Direct) was added for the first 24 h of differentiation/sphere formation. Media changes were performed every 48 h beginning on day 1 of differentiation. On day 7, embryoid bodies were plated to poly-L-ornithine (PLO; Sigma Aldrich, P3655; 20 ng/mL) and natural mouse laminin (Sigma Aldrich, L2020-1MG; 10 µg/mL) coated tissue culture treated dishes. On day 12, rosette structures were manually isolated using a p1000 pipette tip and plated to a new dish. Isolated rosettes were dissociated to single cells with StemPro Accutase (Life Technologies) on day 17 to generate neural stem cells. On d21, neural stem cells were plated for neuronal differentiation in the following: Neurobasal medium, B-27 with vitamin A, GDNF (10 ng/mL), BDNF (10 ng/mL), glutamine (2 mM), and penicillin−streptomycin (50 μg/mL). Neural stem cells were differentiated until day 42 at which point cultures of primarily neurons were isolated for analysis.

Mesodermal differentiation followed published methods with slight modification.^20^ NCRM-5 iPSCs were added to AggreWell 400 plates for embryoid body formation in mesodermal differentiation media consisting of the following: StemPro34 (ThermoFisher), GlutaMAX (2 mM), monothioglycerol (MTG; 4 × 10-4 M, Sigma-Aldrich), penicillin/streptomycin (100 units/mL, ThermoFisher), ascorbic acid (50 µg/ml, Tocris), and BMP-4 (0.5 ng/ml, Peprotech). Y27632 (10 µM) was added for the first 24 h of differentiation/sphere formation. Cells were maintained as floating embryoid bodies (EBs) for 14 days. Growth factors were supplemented to mesodermal differentiation media throughout mesodermal induction. Day 1-4: BMP-4 (10 ng/mL), basic FGF (bFGF; 5 ng/mL, Reprocell) and activin A (3 ng/mL, Peprotech); Day 5–8: VEGF (10 ng/mL, Peprotech) and DKK1 (150 ng/mL, Peprotech); Day 9-12: VEGF (10 ng/mL), DKK1 (150 ng/mL) and bFGF (5 ng/mL). Embryoid bodies were plated to 0.1% gelatin coated dishes on day 15 and cultured for an additional 7 days in mesodermal differentiation media with 4 to 5 biological replicates collected per timepoint. Robust differentiation to neural and mesodermal lineages were confirmed by appropriate cell morphology, including formation of rosette-like structures and neuronal projections during neural differentiation assays and beating-areas and lack of neuronal projections during mesodermal differentiation methods (**Figure 5**). Flash frozen samples were stored at −80 °C until lipid extraction and analysis.

**Figure 5.**
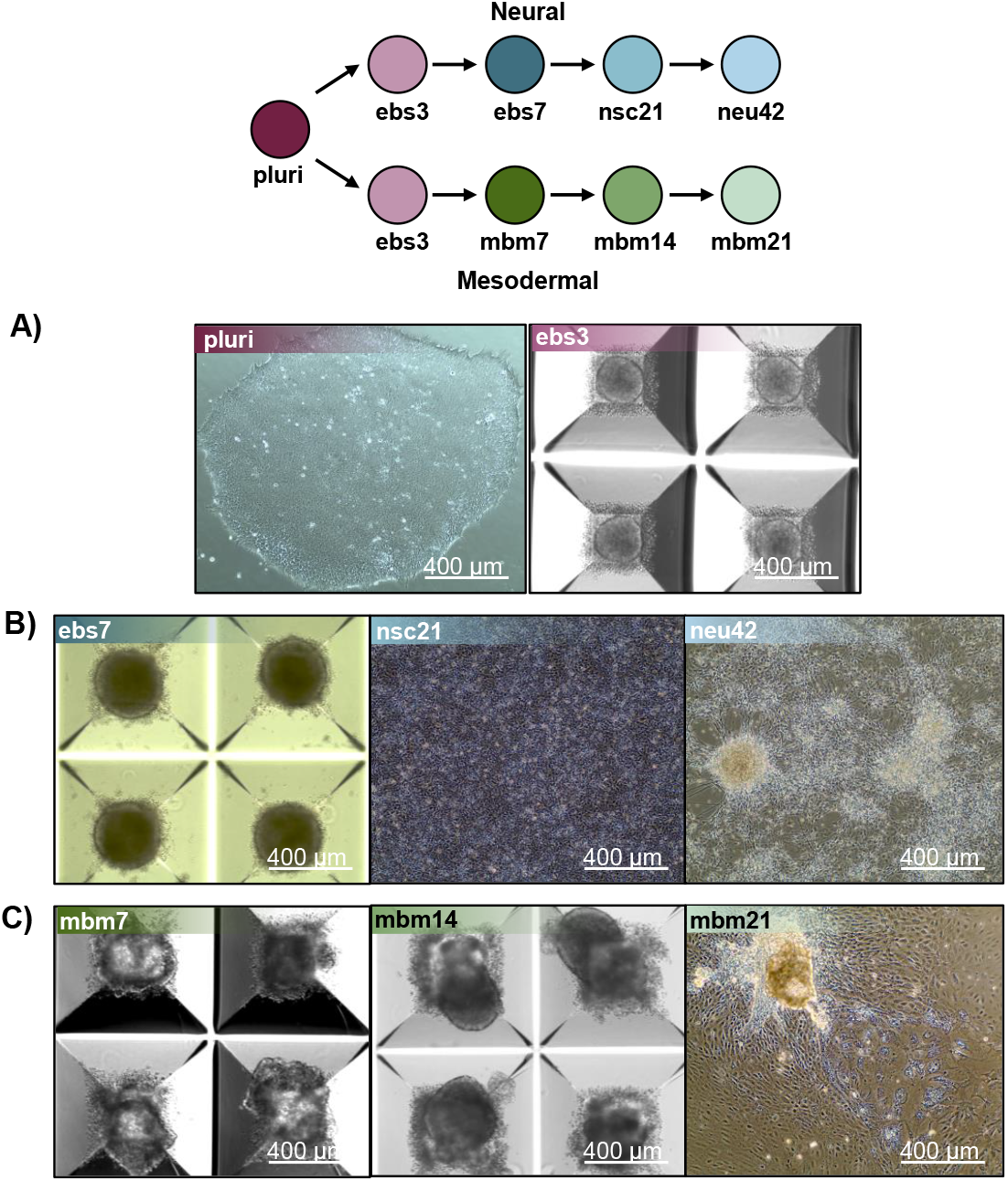
Representative images of iPSC differentiation stages isolated for lipidomics. A) Pluripotent and day 3 embryoid body images undergoing neural induction. B) Unique timepoints of cell lineage images capturing the differentiate of stem cells into neural cells. Timepoints include day 7 neuralized embryoid bodies (ebs7), day 21 neural stem cells (nsc21) and day 42 neurons (neu42). C) Cell images of mesodermal cell differentiation at unique timepoints, including day 7 mesodermal basal medium (mbm7), day 14 mesodermal basal medium (mbm14) and day 21 mesodermal basal medium (mbm21).

#### Lipidomic Extraction

Lipids were extracted following a modified Folch procedure.^50, 51^ To begin, all cell pellets were homogenized with 2.4 mm tungsten beads and 750 µL of −20°C methanol in a bead mill. Samples were subsequently transferred to glass vials and another 750 µL aliquot of cold methanol and 3 mL of chloroform was added. Solutions were incubated for one hour at room temperature. Then, 200 µL of water was added prior to vortexing for 30 s. All samples were then sonicated for 30 min and immediately vortexed for an additional 30 s. Samples were next incubated at 4°C for an hour prior to the addition of a 1.2 mL aliquot of water. All samples were gently mixed then centrifuged for 10 min at 100 x g. From the organic layer of each sample, 300 µL was collected and dried via speedvac. Dried lipid extracts were reconstituted in 10 µL of chloroform and 190 µL methanol and stored at −20°C until LC-IMS-CID-MS analysis.

#### LC-IMS-CID-MS Lipidomic Analysis

Comprehensive lipidomic coverage was accomplished by collecting lipidomic data through four dimensions: liquid chromatography (LC), ion mobility spectrometry (IMS), collision induced dissociation (CID) and mass spectrometry (MS). This was accomplished by coupling an Agilent 1290 Infinity II UHPLC to an Agilent 6560 IMS-MS platform (Santa Clara, CA). A 10 µL injection of each sample was initially separated over a 34 min gradient on a reversed phase Waters CSH column (3.0 mm × 150 mm × 1.7 μm particle size) at a flow rate of 250 μL/min. Detail on gradient ramp, mobile phase composition, and column equilibration is presented in **Table S1**. Eluting lipids were subsequently analyzed using both positive and negative electrospray ionization (ESI) in the 50-1700 m/z range. Lipids were then separated through a DTIMS drift cell containing nitrogen gas at 4 torr.^52^ Finally, ions were fragmented via collision induced dissociation (CID) operating in alternating scans mode to simultaneously collect precursor and product fragment information under a data independent acquisition (DIA) strategy. To optimize CID fragmentation of lipid species, a collision energy ramp was applied based on IMS elution time (**Table S2**).^21, 53^

#### Data Processing, Statistics, and Visualization

The four-dimensional experimental platform leveraged herein results in highly complex data. This study utilized a previously developed Skyline^54^ library of 6,100 transitions for 516 unique lipid molecules published by Kirkwood et al.^55^ to facilitate lipid identifications. Additional data deconvolution software was not explored as data analysis strategies features complementary identification markers including accurate mass, retention time matching, CCS annotation, and product ion fingerprints^.55^ However, lipids with established precedents in the aforementioned cell types but absent within the existing library were enumerated for investigation with LipidCreator.^56^

All annotations were verified by retention time and CCS, additional details are in the Lipid Reporting Checklist. Altogether, a total of 350 lipids were identified in the neural lineage with 177 observed in negative mode, 173 observed in positive mode and 28 of these observed in both modes. Conversely, a total of 453 unique lipids were observed in the mesodermal lineage with a breakdown in observation across negative, positive, and both modes of 269, 184, and 41. Between lineages, an overlap of 170 lipid identifications was observed. All lipid identifications and their respective peak areas were exported for statistical analysis and data visualization in R version 4.0.4.^57^

Outlier assessment, data processing and statistical analysis of data was completed on each stem cell lineage using the pmartR package.^58^ Initially, data was assessed for outliers by testing an RMD-PAV algorithm^59^, Pearson correlation, and PCA clustering. No outliers were observed in any dataset or ionization mode. Data was then normalized by median abundance and log^2^ transformed prior to the statistical analysis of lipid peak areas across subsequent stem cell lineage timepoints. A t-test with an α cut-off of 0.05 and a Holm^60^ multiple comparisons correction was used for all pairwise comparisons. For lipids with missing values, a g-test for qualitative differences across sample types was used to assess statistical significance of events of missingness.^61^ However, the limited sample size of this study resulted in no lipids being returned as statistically significant using this method. Detailed results for both neural and mesodermal stem cell lineages are depicted further within **Tables S3-S4**.

To visualize trends to lipidome alterations overtime in each cell type, significant lipid species were structurally related through head group and FA compositions with the SCOPE cheminformatics toolbox.^23^ Briefly, head group trends were explored through hierarchical clustering of lipids based on the representative SMILES^62^ obtained from the LIPID MAPS^63, 64^ database. Structures were then converted into an ECFP_6^65^ fingerprint and similarity was measured with Tanimoto coefficients and average linkages. FA effects were explored by selectively parsing out lipids based on shared moieties. Given that our analytical method was incapable of differentiating sn-positioning for most of the identified lipids, only FA presence (and not positioning) was considered. Once structural relationships were established, biological results were overlaid by plotting observed log_2_FC of significant lipids using a red/blue gradient to denote up and downregulation. Lipids that were measured but had no statistical significance are depicted as grey. Additional details for the curation of these plots can be found in Odenkirk et al.^23^

## Conclusions

The application of iPSCs remains limited by our understanding of what drives differentiation across cell lineages and facilitates specialized cellular function. This study specifically aimed to evaluate how lipids contribute to the longitudinal development of neural and mesodermal lineages from pluripotent models. A general mechanism of TG and FFA upregulation was observed in the initial neural and mesodermal differentiation timepoints, suggesting common lipid metabolism changes occur upon loss of pluripotency, independent of early lineage commitment. As differentiation continued, distinct phospholipid fluctuations and plasmalogen species changes were observed for both lineages. Specifically, PC O/P and PE O/P lipid upregulation and later downregulation occurred during mesodermal differentiation, while neural differentiation showed a continued increase in PC O/P, LPC and PE O/P lipids. Other unique lineage-specific changes were also observed including for 14:0, 16:0, and 16:1 FAs in neural stem cell differentiation and 20:4 FA up- and subsequent downregulation in both lineages, suggestive of altered eicosanoid intra- and intercellular signalling mechanisms. For the final differentiation timepoints, largely opposite trends were observed among the lipid classes for the neural and mesodermal lineages, demonstrating the uniqueness of lipid composition changes for the two differentiation methods. Overall, these lipidomic analyses demonstrate fluctuations in lipid metabolism during pluripotent stem cell differentiation, detailing basal changes during developmental transitions, and suggesting targeted modulation impacts differentiation and cell type formation. Further studies evaluating how these lipids are driving differentiation and function can now be planned based on these results.

## Supporting information

Lipid_Reporting_Checklist

## Author Information

## Author contributions

MTO, KF, and ESB conceived and designed the research; MTO and KF performed the experiments and analyzed the data; MTO, HCJ, and KF drafted the manuscript; MTO, HCJ, KF, and ESB edited the manuscript. KF and ESB acquired funding. All authors have read and approved the manuscript.

## Data availability

Data for this article, including raw LC-IMS-MS files are available at MassIVE at: https://doi.org/doi:10.25345/C56W96N17. The code for SCOPE analyses using R can be found at: https://github.com/BakerLabNCSU/SCOPE. The version of the code employed for this study is version 4.3.2.

## Acknowledgements

Funding for this work was made possible by grants from the NIH National Institute of Environmental Health Sciences (P42 This journal is © The Royal Society of Chemistry 20xx ES027704), National Institute of General Medical Sciences (R01 GM141277, RM1 GM145416, and P20 GM103620), and a cooperative agreement with the Environmental Protection Agency (STAR RD 84003201). The contents are solely the responsibility of the authors and do not necessarily represent the official views of the funding agencies.

