## Supplementary material for "Lipidomics Reveals Cell Specific Changes During Pluripotent Differentiation to Neural and Mesodermal Lineages": Lipid_Reporting_Checklist

### Contents of Report

Created by https://lipidomicstandards.org, version v2.4.0

**Separation Workflow** 1

### Separation Workflow

#### Overall study design

| | | | |
|---|---|---|---|
| Title of the study | Lipidomics Reveals Cell Specific Changes During Pluripotent Differentiation to Neural and Mesodermal Lineages | | |
| Document creation date | 12/31/2024 | Corresponding Email | |
| Principal investigator | Erin Baker | Is the workflow targeted or untargeted? | Untargeted |
| Institution | University of North Carolina at Chapel Hill | Clinical | No |

#### Lipid extraction

| | | | |
|---|---|---|---|
| Extraction method | 2-phase system | Were internal standards added prior extraction? | No |
| pH adjustment | None | Special conditions | - |
| 2-phase system | Folch | Derivatization | - |

#### Analytical platform

| | | | |
|---|---|---|---|
| Ionizaton additives | Ammonium acetate | MS vendor | Agilent |
| Number of separation dimensions | Two dimensions | Ion source | ESI |
| Separation type 1 | LC | MS Level | MS2 |
| Separation mode 1 (liquid) | RP | Mass window for precursor ion isolation (in Da total isolation window) | 0 |
| Separation window for lipid analyte selection (±) in minutes | 2 | Mass resolution for detected ion at MS2 | High resolution |
| Separation type 2 | IMS | Resolution at m/z 200 at MS2 | 25000 |
| Separation mode 2 (generic) | Drift Tube (N2) | Mass accuracy in ppm at MS2 | 2 |
| Detector | Mass spectrometer | Recording mode of raw data at MS2 | Centroid mode |
| MS type | QTOF | Was/Were additional dimension/techniques used | Yes |

#### Quality control

| | | | |
|---|---|---|---|
| Blanks | Yes | Quality control | Yes |
| Type of Blanks | Extraction blank, Solvent blank | Type of QC sample | Commercial sample |

#### Method qualification and validation

| | |
|---|---|
| Method validation | No |

#### Reporting

| | | | |
|---|---|---|---|
| Are reported raw data uploaded into repository? | Yes | Summary data | Quantification and identification data |
| Link to repository / ID to entry | https://doi.org/doi:10.25345/C5RW06N17 | Raw data upload | Yes |
| Are metadata available? | Yes | Additional comments | all-ions fragmentation was performed after the IMS separation. |

### Sample Descriptions

#### Cell Differentiation / Human / Cells

| | | | |
|---|---|---|---|
| Storage and collection conditions | Available | Additives | None |
| Provided preanalytical information | - | Were samples stored under inert gas? | No |
| Temperature handling original sample | 4-8 °C | Additional preservation methods | No |
| Instant sample preparation | No | Biobank samples | No |
| Storage temperature | -80 °C | | |

### Lipid Class Descriptions

#### 1) CAR, LPC[M+H]+ / Lipid identification

| | | | |
|---|---|---|---|
| Lipid class | CAR, LPC | Limit of detection | No |
| MS Level for identification | MS1, MS2 | RT verified by standard | No |
| Identification level | sn Position | Separation of isobaric/isomeric interferece confirmed | No |
| Polarity mode | Positive | Model for separation prediction | No |
| Type of positive (precursor)ion | [M+H]+ | Additional dimension/techniques | IMS |
| Fragments for identification | Fragment name: (C5H13NO,104) | CCS verified by standard | No |
| Isotope correction at MS1 | No | How was/were the additional dimension(s) used? | For separation of isobaric/isomeric interferece at MS1 and MS2 levels |
| Isotope correction at MS2 | No | Was a model used to predict lipid molecule separation? | No |
| MS1 verified by standard | No | Lipid Identification Software | Skyline |
| MS2 verified by standard | No | Data manipulation | - |
| Background check at MS1 | No | Nomenclature for intact lipid molecule | Yes |
| Background check at MS2 | No | Nomenclature for fragment ions | Yes |
| Did you presume assumptions for identification? | No | Further identification remarks | - |
| Check on: | - | | |

#### 1) CAR, LPC[M+H]+ / Lipid quantification

| | | | |
|---|---|---|---|
| Quantitative | No | Batch correction | No |
| Normalization to reference | No | Further quantification remarks | - |

#### 2) CAR, LPC[M+Na]+ / Lipid identification

| | | | |
|---|---|---|---|
| Lipid class | CAR, LPC | Limit of detection | No |
| MS Level for identification | MS1, MS2 | RT verified by standard | No |
| Identification level | sn Position | Separation of isobaric/isomeric interferece confirmed | No |
| Polarity mode | Positive | Model for separation prediction | No |
| Type of positive (precursor)ion | [M+Na]+ | Additional dimension/techniques | IMS |
| Fragments for identification | | CCS verified by standard | No |

| Fragment name |
|---|
| M+Na-TMA |
| M-HG |

| | | | |
|---|---|---|---|
| Isotope correction at MS1 | No | How was/were the additional dimension(s) used? | For separation of isobaric/isomeric interferece in MS1 and MS2 |
| Isotope correction at MS2 | No | Was a model used to predict lipid molecule separation? | No |
| MS1 verified by standard | No | Lipid Identification Software | Skyline |
| MS2 verified by standard | No | Data manipulation | - |
| Background check at MS1 | No | Nomenclature for intact lipid molecule | Yes |
| Background check at MS2 | No | Nomenclature for fragment ions | N/A |
| Did you presume assumptions for identification? | No | Further identification remarks | - |
| Check on: | - | | |

#### 2) CAR, LPC[M+Na]+ / Lipid quantification

| | | | |
|---|---|---|---|
| Quantitative | No | Batch correction | No |
| Normalization to reference | No | Further quantification remarks | - |

#### 3) CAR, PC[M+H]+ / Lipid identification

| | | | |
|---|---|---|---|
| Lipid class | CAR, PC | Limit of detection | No |
| MS Level for identification | MS1, MS2 | RT verified by standard | Yes |
| Identification level | Molecular species level | Separation of isobaric/isomeric interferece confirmed | No |
| Polarity mode | Positive | Model for separation prediction | No |
| Type of positive (precursor)ion | [M+H]+ | Additional dimension/techniques | IMS |
| Fragments for identification | | CCS verified by standard | Yes |

| Fragment name |
|---|
| M-FA1 |
| M-FA2 |

| | | | |
|---|---|---|---|
| Isotope correction at MS1 | No | How was/were the additional dimension(s) used? | For separation of isobaric/isomeric interferece in MS1 and MS2 dimensions |
| Isotope correction at MS2 | No | Was a model used to predict lipid molecule separation? | No |
| MS1 verified by standard | No | Lipid Identification Software | Skyline |
| MS2 verified by standard | No | Data manipulation | - |
| Background check at MS1 | No | Nomenclature for intact lipid molecule | No |
| Background check at MS2 | No | Nomenclature for fragment ions | N/A |
| Did you presume assumptions for identification? | No | Further identification remarks | - |
| Check on: | - | | |

#### 3) CAR, PC[M+H]+ / Lipid quantification

| | | | |
|---|---|---|---|
| Quantitative | No | Batch correction | No |
| Normalization to reference | No | Further quantification remarks | - |

#### 4) CAR, PC[M+Na]+ / Lipid identification

| | | | |
|---|---|---|---|
| Lipid class | CAR, PC | Limit of detection | No |
| MS Level for identification | MS1, MS2 | RT verified by standard | No |
| Identification level | Molecular species level | Separation of isobaric/isomeric interferece confirmed | No |
| Polarity mode | Positive | Model for separation prediction | No |
| Type of positive (precursor)ion | [M+Na]+ | Additional dimension/techniques | IMS |
| Fragments for identification | | CCS verified by standard | No |

| Fragment name |
|---|
| M+Na-FA1 |
| M+Na-FA2 |
| M+Na-HG |
| M+Na-TMA |
| M+Na-TMA-FA1 |
| M+Na-TMA-FA2 |
| M-FA1 |
| M-FA2 |
| M-HG |
| M-TMA-FA1 |
| M-TMA-FA2 |

| | | | |
|---|---|---|---|
| Isotope correction at MS1 | No | How was/were the additional dimension(s) used? | For separation of isobaric/isomeric interferece in MS1 and MS2 dimensions |
| Isotope correction at MS2 | No | Was a model used to predict lipid molecule separation? | No |
| MS1 verified by standard | No | Lipid Identification Software | Skyline |
| MS2 verified by standard | No | Data manipulation | - |
| Background check at MS1 | No | Nomenclature for intact lipid molecule | No |
| Background check at MS2 | No | Nomenclature for fragment ions | N/A |
| Did you presume assumptions for identification? | No | Further identification remarks | - |
| Check on: | - | | |

#### 4) CAR, PC[M+Na]+ / Lipid quantification

| | | | |
|---|---|---|---|
| Quantitative | No | Batch correction | No |
| Normalization to reference | No | Further quantification remarks | - |

#### 5) CAR, PC P[M+H]+ / Lipid identification

| | | | |
|---|---|---|---|
| Lipid class | CAR, PC P | Limit of detection | No |
| MS Level for identification | MS1, MS2 | RT verified by standard | No |
| Identification level | sn Position | Separation of isobaric/isomeric interferece confirmed | No |
| Polarity mode | Positive | Model for separation prediction | No |
| Type of positive (precursor)ion | [M+H]+ | Additional dimension/techniques | IMS |
| Fragments for identification | | CCS verified by standard | No |

| Fragment name |
|---|
| M-FA1 |
| M-pFA2 |

| | | | |
|---|---|---|---|
| Isotope correction at MS1 | No | How was/were the additional dimension(s) used? | For separation of isobaric/isomeric interferece in MS1 and MS2 dimensions |
| Isotope correction at MS2 | No | Was a model used to predict lipid molecule separation? | No |
| MS1 verified by standard | No | Lipid Identification Software | Skyline |
| MS2 verified by standard | No | Data manipulation | - |
| Background check at MS1 | No | Nomenclature for intact lipid molecule | No |
| Background check at MS2 | No | Nomenclature for fragment ions | N/A |
| Did you presume assumptions for identification? | No | Further identification remarks | - |
| Check on: | - | | |

#### 5) CAR, PC P[M+H]+ / Lipid quantification

| | | | |
|---|---|---|---|
| Quantitative | No | Batch correction | No |
| Normalization to reference | No | Further quantification remarks | - |

#### 6) CAR, PC P[M+Na]+ / Lipid identification

| | | | |
|---|---|---|---|
| Lipid class | CAR, PC P | Limit of detection | No |
| MS Level for identification | MS1, MS2 | RT verified by standard | No |
| Identification level | sn Position | Separation of isobaric/isomeric interferece confirmed | No |
| Polarity mode | Positive | Model for separation prediction | No |
| Type of positive (precursor)ion | [M+Na]+ | Additional dimension/techniques | IMS |
| Fragments for identification | | CCS verified by standard | No |

| Fragment name |
|---|
| M+Na-HG |
| M+Na-TMA |
| M-FA1 |
| M+Na-FA1 |

| | | | |
|---|---|---|---|
| Isotope correction at MS1 | No | How was/were the additional dimension(s) used? | For separation of isobaric/isomeric interferece in MS1 and MS2 dimensions |
| Isotope correction at MS2 | No | Was a model used to predict lipid molecule separation? | No |
| MS1 verified by standard | No | Lipid Identification Software | Skyline |
| MS2 verified by standard | No | Data manipulation | - |
| Background check at MS1 | No | Nomenclature for intact lipid molecule | No |
| Background check at MS2 | No | Nomenclature for fragment ions | N/A |
| Did you presume assumptions for identification? | No | Further identification remarks | - |
| Check on: | - | | |

#### 6) CAR, PC P[M+Na]+ / Lipid quantification

| | | | |
|---|---|---|---|
| Quantitative | No | Batch correction | No |
| Normalization to reference | No | Further quantification remarks | - |

#### 7) CAR, PC[M-H]- / Lipid identification

| | | | |
|---|---|---|---|
| Lipid class | CAR, PC | Limit of detection | No |
| MS Level for identification | MS1, MS2 | RT verified by standard | No |
| Identification level | Molecular species level | Separation of isobaric/isomeric interferece confirmed | No |
| Polarity mode | Negative | Model for separation prediction | No |
| Type of negative (precursor)ion | [M-H]- | Additional dimension/techniques | IMS |
| Fragments for identification | | CCS verified by standard | No |

| Fragment name |
|---|
| -(CH3+CH3COO) |
| -FA1 (+OH) -(CH3+CH3COO) |
| -FA1 (-H) -(CH3+CH3COO) |
| -FA2 (+OH) -(CH3+CH3COO) |
| -FA2 (-H) -(CH3+CH3COO) |
| -FA1 (+O) |
| -FA2 (+O) |
| HG(PC,224) |

| | | | |
|---|---|---|---|
| Isotope correction at MS1 | No | How was/were the additional dimension(s) used? | For separation of isobaric/isomeric interferece in MS1 and MS2 dimensions |
| Isotope correction at MS2 | No | Was a model used to predict lipid molecule separation? | No |
| MS1 verified by standard | No | Lipid Identification Software | Skyline |
| MS2 verified by standard | No | Data manipulation | - |
| Background check at MS1 | No | Nomenclature for intact lipid molecule | No |
| Background check at MS2 | No | Nomenclature for fragment ions | N/A |
| Did you presume assumptions for identification? | No | Further identification remarks | - |
| Check on: | - | | |

#### 7) CAR, PC[M-H]- / Lipid quantification

| | | | |
|---|---|---|---|
| Quantitative | No | Batch correction | No |
| Normalization to reference | No | Further quantification remarks | - |

#### 8) CAR, PG[M-H]- / Lipid identification

| | | | |
|---|---|---|---|
| Lipid class | CAR, PG | Limit of detection | No |
| MS Level for identification | MS1, MS2 | RT verified by standard | No |
| Identification level | Molecular species level | Separation of isobaric/isomeric interferece confirmed | No |
| Polarity mode | Negative | Model for separation prediction | No |
| Type of negative (precursor)ion | [M-H]- | Additional dimension/techniques | IMS |
| Fragments for identification | | CCS verified by standard | No |

| Fragment name |
|---|
| -FA1 (+HO) |
| -FA1 (-H) |
| -FA1(+O) |
| -FA2 (+HO) |
| -FA2 (-H) |
| -FA2(+O) |
| GP(153) |
| HG(PG,171) |
| HG(PG,227) |

| | | | |
|---|---|---|---|
| Isotope correction at MS1 | No | How was/were the additional dimension(s) used? | For separation of isobaric/isomeric interference in MS1 and MS2 dimensions |
| Isotope correction at MS2 | No | Was a model used to predict lipid molecule separation? | No |
| MS1 verified by standard | No | Lipid Identification Software | Skyline |
| MS2 verified by standard | No | Data manipulation | - |
| Background check at MS1 | No | Nomenclature for intact lipid molecule | No |
| Background check at MS2 | No | Nomenclature for fragment ions | N/A |
| Did you presume assumptions for identification? | No | Further identification remarks | - |
| Check on: | - | | |

#### 8) CAR, PG[M-H]- / Lipid quantification

| | | | |
|---|---|---|---|
| Quantitative | No | Batch correction | No |
| Normalization to reference | No | Further quantification remarks | - |

#### 9) CAR, PS[M-H]- / Lipid identification

| | | | |
|---|---|---|---|
| Lipid class | CAR, PS | Limit of detection | No |
| MS Level for identification | MS1, MS2 | RT verified by standard | No |
| Identification level | Molecular species level | Separation of isobaric/isomeric interferece confirmed | No |
| Polarity mode | Negative | Model for separation prediction | No |
| Type of negative (precursor)ion | [M-H]- | Additional dimension/techniques | IMS |
| Fragments for identification | | CCS verified by standard | No |

| Fragment name |
|---|
| -(C3H5NO2,87) |
| -FA1 (+OH) |
| -FA1 (-H) |
| -FA2 (+OH) |
| -FA2 (-H) |
| -FA1 (+O) |
| -FA2 (+O) |
| GP(153) |

| | | | |
|---|---|---|---|
| Isotope correction at MS1 | No | How was/were the additional dimension(s) used? | For separation of isobaric/isomeric interferece in MS1 and MS2 dimensions |
| Isotope correction at MS2 | No | Was a model used to predict lipid molecule separation? | No |
| MS1 verified by standard | No | Lipid Identification Software | Skyline |
| MS2 verified by standard | No | Data manipulation | - |
| Background check at MS1 | No | Nomenclature for intact lipid molecule | No |
| Background check at MS2 | No | Nomenclature for fragment ions | N/A |
| Did you presume assumptions for identification? | No | Further identification remarks | - |
| Check on: | - | | |

#### 9) CAR, PS[M-H]- / Lipid quantification

| | | | |
|---|---|---|---|
| Quantitative | No | Batch correction | No |
| Normalization to reference | No | Further quantification remarks | - |

#### 10) CAR, PE[M-H]- / Lipid identification

| | | | |
|---|---|---|---|
| Lipid class | CAR, PE | Limit of detection | No |
| MS Level for identification | MS1, MS2 | RT verified by standard | No |
| Identification level | Molecular species level | Separation of isobaric/isomeric interferece confirmed | No |
| Polarity mode | Negative | Model for separation prediction | No |
| Type of negative (precursor)ion | [M-H]- | Additional dimension/techniques | IMS |
| Fragments for identification | | CCS verified by standard | No |

| Fragment name |
|---|
| -FA1 (+OH) |
| -FA1 (+O) |
| -FA1 (-H) |
| -FA2 (+OH) |
| -FA2 (+O) |
| -FA2 (-H) |
| GP(153) |
| HG(PE,196) |

| | | | |
|---|---|---|---|
| Isotope correction at MS1 | No | How was/were the additional dimension(s) used? | For separation of isobaric/isomeric interferece in MS1 and MS2 dimensions |
| Isotope correction at MS2 | No | Was a model used to predict lipid molecule separation? | No |
| MS1 verified by standard | No | Lipid Identification Software | Skyline |
| MS2 verified by standard | No | Data manipulation | - |
| Background check at MS1 | No | Nomenclature for intact lipid molecule | No |
| Background check at MS2 | No | Nomenclature for fragment ions | N/A |
| Did you presume assumptions for identification? | No | Further identification remarks | - |
| Check on: | - | | |

#### 10) CAR, PE[M-H]- / Lipid quantification

| | | | |
|---|---|---|---|
| Quantitative | No | Batch correction | No |
| Normalization to reference | No | Further quantification remarks | - |

#### 11) CAR, Cer[M+CH3COO]- / Lipid identification

| | | | |
|---|---|---|---|
| Lipid class | CAR, Cer | Limit of detection | No |
| MS Level for identification | MS1, MS2 | RT verified by standard | No |
| Identification level | Species level | Separation of isobaric/isomeric interferece confirmed | No |
| Polarity mode | Negative | Model for separation prediction | No |
| Type of negative (precursor)ion | [M+CH3COO]- | Additional dimension/techniques | IMS |
| Fragments for identification | | CCS verified by standard | No |

| Fragment name |
|---|
| FA (+C2H3N) |
| FA (+C2H3NO) |
| FA (+HN) |
| LCB (-C2H8NO) |
| LCB (-CH3O) |
| LCB (-H6NO) |

| | | | |
|---|---|---|---|
| Isotope correction at MS1 | No | How was/were the additional dimension(s) used? | For separation of isobaric/isomeric interferece in MS1 and MS2 dimensions |
| Isotope correction at MS2 | No | Was a model used to predict lipid molecule separation? | No |
| MS1 verified by standard | No | Lipid Identification Software | Skyline |
| MS2 verified by standard | No | Data manipulation | - |
| Background check at MS1 | No | Nomenclature for intact lipid molecule | No |
| Background check at MS2 | No | Nomenclature for fragment ions | N/A |
| Did you presume assumptions for identification? | No | Further identification remarks | - |
| Check on: | - | | |

#### 11) CAR, Cer[M+CH3COO]- / Lipid quantification

| | | | |
|---|---|---|---|
| Quantitative | No | Batch correction | No |
| Normalization to reference | No | Further quantification remarks | - |

#### 12) CAR, Cer[M+HCOO]- / Lipid identification

| | | | |
|---|---|---|---|
| Lipid class | CAR, Cer | Limit of detection | No |
| MS Level for identification | MS1, MS2 | RT verified by standard | No |
| Identification level | Molecular species level | Separation of isobaric/isomeric interferece confirmed | No |
| Polarity mode | Negative | Model for separation prediction | No |
| Type of negative (precursor)ion | [M+HCOO]- | Additional dimension/techniques | IMS |
| Fragments for identification | | CCS verified by standard | No |

| Fragment name |
|---|
| FA (+C2H3N) |
| FA (+C2H3NO) |
| FA (+HN) |
| LCB (-C2H8NO) |
| LCB (-CH3O) |
| LCB (-H6NO) |

| | | | |
|---|---|---|---|
| Isotope correction at MS1 | No | How was/were the additional dimension(s) used? | For separation of isobaric/isomeric interferece in MS1 and MS2 dimensions |
| Isotope correction at MS2 | No | Was a model used to predict lipid molecule separation? | No |
| MS1 verified by standard | No | Lipid Identification Software | Skyline |
| MS2 verified by standard | No | Data manipulation | - |
| Background check at MS1 | No | Nomenclature for intact lipid molecule | No |
| Background check at MS2 | No | Nomenclature for fragment ions | N/A |
| Did you presume assumptions for identification? | No | Further identification remarks | - |
| Check on: | - | | |

#### 12) CAR, Cer[M+HCOO]- / Lipid quantification

| | | | |
|---|---|---|---|
| Quantitative | No | Batch correction | No |
| Normalization to reference | No | Further quantification remarks | - |

#### 13) CAR, Cer[M-H]- / Lipid identification

| | | | |
|---|---|---|---|
| Lipid class | CAR, Cer | Limit of detection | No |
| MS Level for identification | MS1, MS2 | RT verified by standard | No |
| Identification level | Molecular species level | Separation of isobaric/isomeric interferece confirmed | No |
| Polarity mode | Negative | Model for separation prediction | No |
| Type of negative (precursor)ion | [M-H]- | Additional dimension/techniques | IMS |
| Fragments for identification | | CCS verified by standard | No |

| Fragment name |
|---|
| FA (+C2H3N) |
| FA (+C2H3NO) |
| FA (+HN) |
| LCB (-C2H8NO) |
| LCB (-CH3O) |
| LCB (-H6NO) |

| | | | |
|---|---|---|---|
| Isotope correction at MS1 | No | How was/were the additional dimension(s) used? | For separation of isobaric/isomeric interferece in MS1 and MS2 dimensions |
| Isotope correction at MS2 | No | Was a model used to predict lipid molecule separation? | No |
| MS1 verified by standard | No | Lipid Identification Software | Skyline |
| MS2 verified by standard | No | Data manipulation | - |
| Background check at MS1 | No | Nomenclature for intact lipid molecule | No |
| Background check at MS2 | No | Nomenclature for fragment ions | N/A |
| Did you presume assumptions for identification? | No | Further identification remarks | - |
| Check on: | - | | |

#### 13) CAR, Cer[M-H]- / Lipid quantification

| | | | |
|---|---|---|---|
| Quantitative | No | Batch correction | No |
| Normalization to reference | No | Further quantification remarks | - |

#### 14) CAR, FA[M-H]- / Lipid identification

| | | | |
|---|---|---|---|
| Lipid class | CAR, FA | RT verified by standard | No |
| MS Level for identification | MS1 | Separation of isobaric/isomeric interferece confirmed | No |
| Identification level | Species level | Model for separation prediction | No |
| Polarity mode | Negative | Additional dimension/techniques | IMS |
| Type of negative (precursor)ion | [M-H]- | CCS verified by standard | No |
| Isotope correction at MS1 | No | How was/were the additional dimension(s) used? | For separation of isobaric/isomeric interferece in MS1 and MS2 dimensions |
| MS1 verified by standard | No | Was a model used to predict lipid molecule separation? | No |
| Background check at MS1 | No | Lipid Identification Software | Skyline |
| Did you presume assumptions for identification? | No | Data manipulation | - |
| Check on: | - | Nomenclature for intact lipid molecule | No |
| Limit of detection | No | Further identification remarks | - |

#### 14) CAR, FA[M-H]- / Lipid quantification

| | | | |
|---|---|---|---|
| Quantitative | No | Batch correction | No |
| Normalization to reference | No | Further quantification remarks | - |

#### 15) CAR, GM3[M-H]- / Lipid identification

| | | | |
|---|---|---|---|
| Lipid class | CAR, GM3 | Limit of detection | No |
| MS Level for identification | MS1, MS2 | RT verified by standard | No |
| Identification level | Species level | Separation of isobaric/isomeric interferece confirmed | No |
| Polarity mode | Negative | Model for separation prediction | No |
| Type of negative (precursor)ion | [M-H]- | Additional dimension/techniques | IMS |
| Fragments for identification | | CCS verified by standard | No |

| Fragment name |
|---|
| -HG(NHex,291) |
| -HG(NHex2,453) |
| -HG(NHex3,615) |
| -HG(NHex2,471) |
| -HG(NHex2,633) |
| HG(NHex, 290) |

| | | | |
|---|---|---|---|
| Isotope correction at MS1 | No | How was/were the additional dimension(s) used? | For separation of isobaric/isomeric interferece in MS1 and Ms2 dimensions |
| Isotope correction at MS2 | No | Was a model used to predict lipid molecule separation? | No |
| MS1 verified by standard | No | Lipid Identification Software | Skyline |
| MS2 verified by standard | No | Data manipulation | - |
| Background check at MS1 | No | Nomenclature for intact lipid molecule | No |
| Background check at MS2 | No | Nomenclature for fragment ions | N/A |
| Did you presume assumptions for identification? | No | Further identification remarks | - |
| Check on: | - | | |

#### 15) CAR, GM3[M-H]- / Lipid quantification

| | | | |
|---|---|---|---|
| Quantitative | No | Batch correction | No |
| Normalization to reference | No | Further quantification remarks | - |

#### 16) CAR, HexCer[M+CH3COO]- / Lipid identification

| | | | |
|---|---|---|---|
| Lipid class | CAR, HexCer | Limit of detection | No |
| MS Level for identification | MS1, MS2 | RT verified by standard | No |
| Identification level | Molecular species level | Separation of isobaric/isomeric interferece confirmed | No |
| Polarity mode | Negative | Model for separation prediction | No |
| Type of negative (precursor)ion | [M+CH3COO]- | Additional dimension/techniques | IMS |
| Fragments for identification | | CCS verified by standard | No |

| Fragment name |
|---|
| -HG(Hex,162) |
| -HG(Hex,180) |
| FA (+C2H3N) |
| FA (+C2H3NO) |
| FA (+NO) |
| LCB (-CH3O) |
| LCB (-H6NO) |
| LCB (-C2H8NO) |

| | | | |
|---|---|---|---|
| Isotope correction at MS1 | No | How was/were the additional dimension(s) used? | For separation of isobaric/isomeric interferece of MS1 and MS2 dimensions |
| Isotope correction at MS2 | No | Was a model used to predict lipid molecule separation? | No |
| MS1 verified by standard | No | Lipid Identification Software | Skyline |
| MS2 verified by standard | No | Data manipulation | - |
| Background check at MS1 | No | Nomenclature for intact lipid molecule | No |
| Background check at MS2 | No | Nomenclature for fragment ions | N/A |
| Did you presume assumptions for identification? | No | Further identification remarks | - |
| Check on: | - | | |

#### 16) CAR, HexCer[M+CH3COO]- / Lipid quantification

| | | | |
|---|---|---|---|
| Quantitative | No | Batch correction | No |
| Normalization to reference | No | Further quantification remarks | - |

#### 17) CAR, HexCer[M+HCOO]- / Lipid identification

| | | | |
|---|---|---|---|
| Lipid class | CAR, HexCer | Limit of detection | No |
| MS Level for identification | MS1, MS2 | RT verified by standard | No |
| Identification level | Molecular species level | Separation of isobaric/isomeric interferece confirmed | No |
| Polarity mode | Negative | Model for separation prediction | No |
| Type of negative (precursor)ion | [M+HCOO]- | Additional dimension/techniques | IMS |
| Fragments for identification | | CCS verified by standard | No |

| Fragment name |
|---|
| -HG(Hex,162) |
| -HG(Hex,180) |
| FA (+C2H3N) |
| FA (+C2H3NO) |
| FA (+HN) |
| LCB (-C2H8NO) |
| LCB (-CH3NO) |
| LCB (-H6NO) |

| | | | |
|---|---|---|---|
| Isotope correction at MS1 | No | How was/were the additional dimension(s) used? | For separation of isobaric/isomeric interferece of MS1 and MS2 dimensions |
| Isotope correction at MS2 | No | Was a model used to predict lipid molecule separation? | No |
| MS1 verified by standard | No | Lipid Identification Software | Skyline |
| MS2 verified by standard | No | Data manipulation | - |
| Background check at MS1 | No | Nomenclature for intact lipid molecule | No |
| Background check at MS2 | No | Nomenclature for fragment ions | N/A |
| Did you presume assumptions for identification? | No | Further identification remarks | - |
| Check on: | - | | |

#### 17) CAR, HexCer[M+HCOO]- / Lipid quantification

| | | | |
|---|---|---|---|
| Quantitative | No | Batch correction | No |
| Normalization to reference | No | Further quantification remarks | - |

#### 18) CAR, HexCer[M-H]- / Lipid identification

| | | | |
|---|---|---|---|
| Lipid class | CAR, HexCer | Limit of detection | No |
| MS Level for identification | MS1, MS2 | RT verified by standard | No |
| Identification level | Molecular species level | Separation of isobaric/isomeric interferece confirmed | No |
| Polarity mode | Negative | Model for separation prediction | No |
| Type of negative (precursor)ion | [M-H]- | Additional dimension/techniques | IMS |
| Fragments for identification | | CCS verified by standard | No |

| Fragment name |
|---|
| FA (+C2H3O) |
| FA (+C2H3NO) |
| FA (+HN) |
| LCB (-C2H8NO) |
| LCB (-CH3O) |
| LCB (-H6NO) |

| | | | |
|---|---|---|---|
| Isotope correction at MS1 | No | How was/were the additional dimension(s) used? | For separation of isobaric/isomeric interferece of MS1 and MS2 dimensions |
| Isotope correction at MS2 | No | Was a model used to predict lipid molecule separation? | No |
| MS1 verified by standard | No | Lipid Identification Software | Skyline |
| MS2 verified by standard | No | Data manipulation | - |
| Background check at MS1 | No | Nomenclature for intact lipid molecule | No |
| Background check at MS2 | No | Nomenclature for fragment ions | N/A |
| Did you presume assumptions for identification? | No | Further identification remarks | - |
| Check on: | - | | |

#### 18) CAR, HexCer[M-H]- / Lipid quantification

| | | | |
|---|---|---|---|
| Quantitative | No | Batch correction | No |
| Normalization to reference | No | Further quantification remarks | - |

#### 19) CAR, LPC[M+CH3COO]- / Lipid identification

| | | | |
|---|---|---|---|
| Lipid class | CAR, LPC | Limit of detection | No |
| MS Level for identification | MS1, MS2 | RT verified by standard | No |
| Identification level | sn Position | Separation of isobaric/isomeric interferece confirmed | No |
| Polarity mode | Negative | Model for separation prediction | No |
| Type of negative (precursor)ion | [M+CH3COO]- | Additional dimension/techniques | IMS |
| Fragments for identification | | CCS verified by standard | No |

| Fragment name |
|---|
| HG(PC,224) |
| FA1(+O) |
| -(CH3+CH3COO) |

| | | | |
|---|---|---|---|
| Isotope correction at MS1 | No | How was/were the additional dimension(s) used? | For separation of isobaric/isomeric interferece of MS1 and MS2 dimensions |
| Isotope correction at MS2 | No | Was a model used to predict lipid molecule separation? | No |
| MS1 verified by standard | No | Lipid Identification Software | Skyline |
| MS2 verified by standard | No | Data manipulation | - |
| Background check at MS1 | No | Nomenclature for intact lipid molecule | No |
| Background check at MS2 | No | Nomenclature for fragment ions | N/A |
| Did you presume assumptions for identification? | No | Further identification remarks | - |
| Check on: | - | | |

#### 19) CAR, LPC[M+CH3COO]- / Lipid quantification

| | | | |
|---|---|---|---|
| Quantitative | No | Batch correction | No |
| Normalization to reference | No | Further quantification remarks | - |

#### 20) CAR, LPE[M-H]- / Lipid identification

| | | | |
|---|---|---|---|
| Lipid class | CAR, LPE | Limit of detection | No |
| MS Level for identification | MS1, MS2 | RT verified by standard | No |
| Identification level | sn Position | Separation of isobaric/isomeric interferece confirmed | No |
| Polarity mode | Negative | Model for separation prediction | No |
| Type of negative (precursor)ion | [M-H]- | Additional dimension/techniques | IMS |
| Fragments for identification | | CCS verified by standard | No |

| Fragment name |
|---|
| -FA1(-H)-(H2O) |
| -FA1(-H) |
| GP(153) |
| -FA1(+O) |

| | | | |
|---|---|---|---|
| Isotope correction at MS1 | No | How was/were the additional dimension(s) used? | For separation of isobaric/isomeric interferece for MS1 and MS2 dimensions |
| Isotope correction at MS2 | No | Was a model used to predict lipid molecule separation? | No |
| MS1 verified by standard | No | Lipid Identification Software | Skyline |
| MS2 verified by standard | No | Data manipulation | - |
| Background check at MS1 | No | Nomenclature for intact lipid molecule | No |
| Background check at MS2 | No | Nomenclature for fragment ions | N/A |
| Did you presume assumptions for identification? | No | Further identification remarks | - |
| Check on: | - | | |

#### 20) CAR, LPE[M-H]- / Lipid quantification

| | | | |
|---|---|---|---|
| Quantitative | No | Batch correction | No |
| Normalization to reference | No | Further quantification remarks | - |

#### 21) CAR, LPA[M-H]- / Lipid identification

| | | | |
|---|---|---|---|
| Lipid class | CAR, LPA | Limit of detection | No |
| MS Level for identification | MS1, MS2 | RT verified by standard | No |
| Identification level | Molecular species level | Separation of isobaric/isomeric interferece confirmed | No |
| Polarity mode | Negative | Model for separation prediction | No |
| Type of negative (precursor)ion | [M-H]- | Additional dimension/techniques | IMS |
| Fragments for identification | | CCS verified by standard | No |

| Fragment name |
|---|
| GP(153) |
| P(79) |
| FA1 (+O) |

| | | | |
|---|---|---|---|
| Isotope correction at MS1 | No | How was/were the additional dimension(s) used? | For separation of isobaric/isomeric interferece for MS1 and MS2 dimensions |
| Isotope correction at MS2 | No | Was a model used to predict lipid molecule separation? | No |
| MS1 verified by standard | No | Lipid Identification Software | Skyline |
| MS2 verified by standard | No | Data manipulation | - |
| Background check at MS1 | No | Nomenclature for intact lipid molecule | No |
| Background check at MS2 | No | Nomenclature for fragment ions | N/A |
| Did you presume assumptions for identification? | No | Further identification remarks | - |
| Check on: | - | | |

#### 21) CAR, LPA[M-H]- / Lipid quantification

| | | | |
|---|---|---|---|
| Quantitative | No | Batch correction | No |
| Normalization to reference | No | Further quantification remarks | - |

#### 22) CAR, LPG[M-H]- / Lipid identification

| | | | |
|---|---|---|---|
| Lipid class | CAR, LPG | Limit of detection | No |
| MS Level for identification | MS1, MS2 | RT verified by standard | No |
| Identification level | Molecular species level | Separation of isobaric/isomeric interferece confirmed | No |
| Polarity mode | Negative | Model for separation prediction | No |
| Type of negative (precursor)ion | [M-H]- | Additional dimension/techniques | IMS |
| Fragments for identification | | CCS verified by standard | No |

| Fragment name |
|---|
| GP(153) |
| -FA1(-H) |
| -FA1(+HO) |
| -FA1(+O) |

| | | | |
|---|---|---|---|
| Isotope correction at MS1 | No | How was/were the additional dimension(s) used? | For separation of isobaric/isomeric interferece for MS1 and MS2 dimensions |
| Isotope correction at MS2 | No | Was a model used to predict lipid molecule separation? | No |
| MS1 verified by standard | No | Lipid Identification Software | Skyline |
| MS2 verified by standard | No | Data manipulation | - |
| Background check at MS1 | No | Nomenclature for intact lipid molecule | No |
| Background check at MS2 | No | Nomenclature for fragment ions | N/A |
| Did you presume assumptions for identification? | No | Further identification remarks | - |
| Check on: | - | | |

#### 22) CAR, LPG[M-H]- / Lipid quantification

| | | | |
|---|---|---|---|
| Quantitative | No | Batch correction | No |
| Normalization to reference | No | Further quantification remarks | - |

#### 23) CAR, PA[M-H]- / Lipid identification

| | | | |
|---|---|---|---|
| Lipid class | CAR, PA | Limit of detection | No |
| MS Level for identification | MS1, MS2 | RT verified by standard | No |
| Identification level | Molecular species level | Separation of isobaric/isomeric interferece confirmed | No |
| Polarity mode | Negative | Model for separation prediction | No |
| Type of negative (precursor)ion | [M-H]- | Additional dimension/techniques | IMS |
| Fragments for identification | | CCS verified by standard | No |

| Fragment name |
|---|
| GP(153) |
| FA1(+O) |
| FA1(+HO) |
| FA1(-H) |
| FA2(+O) |
| FA2(+HO) |
| FA2(-H) |

| | | | |
|---|---|---|---|
| Isotope correction at MS1 | No | How was/were the additional dimension(s) used? | For separation of isobaric/isomeric interferece for MS1 and MS2 dimensions |
| Isotope correction at MS2 | No | Was a model used to predict lipid molecule separation? | No |
| MS1 verified by standard | No | Lipid Identification Software | Skyline |
| MS2 verified by standard | No | Data manipulation | - |
| Background check at MS1 | No | Nomenclature for intact lipid molecule | No |
| Background check at MS2 | No | Nomenclature for fragment ions | N/A |
| Did you presume assumptions for identification? | No | Further identification remarks | - |
| Check on: | - | | |

#### 23) CAR, PA[M-H]- / Lipid quantification

| | | | |
|---|---|---|---|
| Quantitative | No | Batch correction | No |
| Normalization to reference | No | Further quantification remarks | - |

#### 24) CAR, PC P[M+CH3COO]- / Lipid identification

| | | | |
|---|---|---|---|
| Lipid class | CAR, PC P | Limit of detection | No |
| MS Level for identification | MS1, MS2 | RT verified by standard | No |
| Identification level | Molecular species level | Separation of isobaric/isomeric interferece confirmed | No |
| Polarity mode | Negative | Model for separation prediction | No |
| Type of negative (precursor)ion | [M+CH3COO]- | Additional dimension/techniques | IMS |
| Fragments for identification | | CCS verified by standard | No |

| Fragment name |
|---|
| HG(PC)-(CH3+CH3COO) |
| (CH3+CH3COO) |
| FA1 (+OH) |
| FA1 (-CO) |
| FA1 (+O) |
| FA1 (-H) |
| FA O-[xx:x] |

| | | | |
|---|---|---|---|
| Isotope correction at MS1 | No | How was/were the additional dimension(s) used? | For separation of isobaric/isomeric interferece for MS1 and MS2 identifications |
| Isotope correction at MS2 | No | Was a model used to predict lipid molecule separation? | No |
| MS1 verified by standard | No | Lipid Identification Software | Skyline |
| MS2 verified by standard | No | Data manipulation | - |
| Background check at MS1 | No | Nomenclature for intact lipid molecule | No |
| Background check at MS2 | No | Nomenclature for fragment ions | N/A |
| Did you presume assumptions for identification? | No | Further identification remarks | - |
| Check on: | - | | |

#### 24) CAR, PC P[M+CH3COO]- / Lipid quantification

| | | | |
|---|---|---|---|
| Quantitative | No | Batch correction | No |
| Normalization to reference | No | Further quantification remarks | - |

#### 25) CAR, PC O[M+CH3COO]- / Lipid identification

| | | | |
|---|---|---|---|
| Lipid class | CAR, PC O | Limit of detection | No |
| MS Level for identification | MS1, MS2 | RT verified by standard | No |
| Identification level | Molecular species level | Separation of isobaric/isomeric interferece confirmed | No |
| Polarity mode | Negative | Model for separation prediction | No |
| Type of negative (precursor)ion | [M+CH3COO]- | Additional dimension/techniques | IMS |
| Fragments for identification | | CCS verified by standard | No |

| Fragment name |
|---|
| HG(PC)-(CH3+CH3COO) |
| FA O-[xx:x] |
| FA1(+HO) |
| FA1(+O) |
| FA1-(CO) |
| FA1(-H) |
| -(CH3+CH3COO) |

| | | | |
|---|---|---|---|
| Isotope correction at MS1 | No | How was/were the additional dimension(s) used? | For separation of isobaric/isomeric interferece for MS1 and MS2 dimensions |
| Isotope correction at MS2 | No | Was a model used to predict lipid molecule separation? | No |
| MS1 verified by standard | No | Lipid Identification Software | Skyline |
| MS2 verified by standard | No | Data manipulation | - |
| Background check at MS1 | No | Nomenclature for intact lipid molecule | No |
| Background check at MS2 | No | Nomenclature for fragment ions | N/A |
| Did you presume assumptions for identification? | No | Further identification remarks | - |
| Check on: | - | | |

#### 25) CAR, PC O[M+CH3COO]- / Lipid quantification

| | | | |
|---|---|---|---|
| Quantitative | No | Batch correction | No |
| Normalization to reference | No | Further quantification remarks | - |

#### 26) CAR, PI[M-H]- / Lipid identification

| | | | |
|---|---|---|---|
| Lipid class | CAR, PI | Limit of detection | No |
| MS Level for identification | MS1, MS2 | RT verified by standard | No |
| Identification level | Molecular species level | Separation of isobaric/isomeric interferece confirmed | No |
| Polarity mode | Negative | Model for separation prediction | No |
| Type of negative (precursor)ion | [M-H]- | Additional dimension/techniques | IMS |
| Fragments for identification | | CCS verified by standard | No |

| Fragment name |
|---|
| -FA1(-H) |
| FA1(+O) |
| -FA1(+HO) |
| -FA2(-H) |
| -FA2(+O) |
| -FA2(+HO) |
| GP(153) |
| HG(PI,241) |

| | | | |
|---|---|---|---|
| Isotope correction at MS1 | No | How was/were the additional dimension(s) used? | For separation of isobaric/isomeric interferece in MS1 and MS2 dimensions |
| Isotope correction at MS2 | No | Was a model used to predict lipid molecule separation? | No |
| MS1 verified by standard | No | Lipid Identification Software | Skyline |
| MS2 verified by standard | No | Data manipulation | - |
| Background check at MS1 | No | Nomenclature for intact lipid molecule | No |
| Background check at MS2 | No | Nomenclature for fragment ions | N/A |
| Did you presume assumptions for identification? | No | Further identification remarks | - |
| Check on: | - | | |

#### 26) CAR, PI[M-H]- / Lipid quantification

| | | | |
|---|---|---|---|
| Quantitative | No | Batch correction | No |
| Normalization to reference | No | Further quantification remarks | - |

#### 27) CAR, PE P[M-H]- / Lipid identification

| | | | |
|---|---|---|---|
| Lipid class | CAR, PE P | Limit of detection | No |
| MS Level for identification | MS1, MS2 | RT verified by standard | No |
| Identification level | Molecular species level | Separation of isobaric/isomeric interferece confirmed | No |
| Polarity mode | Negative | Model for separation prediction | No |
| Type of negative (precursor)ion | [M-H]- | Additional dimension/techniques | IMS |
| Fragments for identification | | CCS verified by standard | No |

| Fragment name |
|---|
| FA2 -(CO) |
| -FA2(-H) |
| -FA2(+HO) |
| FA2(+O) |
| FA O-[xx:x] |
| HG(PE,196) |

| | | | |
|---|---|---|---|
| Isotope correction at MS1 | No | How was/were the additional dimension(s) used? | For separation of isobaric/isomeric interferece in MS1 and MS21 dimensions |
| Isotope correction at MS2 | No | Was a model used to predict lipid molecule separation? | No |
| MS1 verified by standard | No | Lipid Identification Software | Skyline |
| MS2 verified by standard | No | Data manipulation | - |
| Background check at MS1 | No | Nomenclature for intact lipid molecule | No |
| Background check at MS2 | No | Nomenclature for fragment ions | N/A |
| Did you presume assumptions for identification? | No | Further identification remarks | - |
| Check on: | - | | |

#### 27) CAR, PE P[M-H]- / Lipid quantification

| | | | |
|---|---|---|---|
| Quantitative | No | Batch correction | No |
| Normalization to reference | No | Further quantification remarks | - |

#### 28) CAR, PE O[M-H]- / Lipid identification

| | | | |
|---|---|---|---|
| Lipid class | CAR, PE O | Limit of detection | No |
| MS Level for identification | MS1, MS2 | RT verified by standard | No |
| Identification level | Molecular species level | Separation of isobaric/isomeric interferece confirmed | No |
| Polarity mode | Negative | Model for separation prediction | No |
| Type of negative (precursor)ion | [M-H]- | Additional dimension/techniques | IMS |
| Fragments for identification | | CCS verified by standard | No |

| Fragment name |
|---|
| FA2 -(CO) |
| -FA2(-H) |
| -FA2(+HO) |
| FA2(+O) |
| GP(135) |
| GP(153) |

| | | | |
|---|---|---|---|
| Isotope correction at MS1 | No | How was/were the additional dimension(s) used? | For separation of isobaric/isomeric interferece in MS1 and MS2 dimensions |
| Isotope correction at MS2 | No | Was a model used to predict lipid molecule separation? | No |
| MS1 verified by standard | No | Lipid Identification Software | Skyline |
| MS2 verified by standard | No | Data manipulation | - |
| Background check at MS1 | No | Nomenclature for intact lipid molecule | No |
| Background check at MS2 | No | Nomenclature for fragment ions | N/A |
| Did you presume assumptions for identification? | No | Further identification remarks | - |
| Check on: | - | | |

#### 28) CAR, PE O[M-H]- / Lipid quantification

| | | | |
|---|---|---|---|
| Quantitative | No | Batch correction | No |
| Normalization to reference | No | Further quantification remarks | - |

#### 29) CAR, SM[M-H]- / Lipid identification

| | | | |
|---|---|---|---|
| Lipid class | CAR, SM | Limit of detection | No |
| MS Level for identification | MS1, MS2 | RT verified by standard | No |
| Identification level | Molecular species level | Separation of isobaric/isomeric interferece confirmed | No |
| Polarity mode | Negative | Model for separation prediction | No |
| Type of negative (precursor)ion | [M-H]- | Additional dimension/techniques | IMS |
| Fragments for identification | | CCS verified by standard | No |

| Fragment name |
|---|
| HG(PC,168) |
| FA1(+O) |

| | | | |
|---|---|---|---|
| Isotope correction at MS1 | No | How was/were the additional dimension(s) used? | For separation of isobaric/isomeric interferece in MS1 and MS2 dimensions |
| Isotope correction at MS2 | No | Was a model used to predict lipid molecule separation? | No |
| MS1 verified by standard | No | Lipid Identification Software | Skyline |
| MS2 verified by standard | No | Data manipulation | - |
| Background check at MS1 | No | Nomenclature for intact lipid molecule | No |
| Background check at MS2 | No | Nomenclature for fragment ions | N/A |
| Did you presume assumptions for identification? | No | Further identification remarks | - |
| Check on: | - | | |

#### 29) CAR, SM[M-H]- / Lipid quantification

| | | | |
|---|---|---|---|
| Quantitative | No | Batch correction | No |
| Normalization to reference | No | Further quantification remarks | - |

#### 30) CAR, AC[M+H]+ / Lipid identification

| | | | |
|---|---|---|---|
| Lipid class | CAR, AC | Limit of detection | No |
| MS Level for identification | MS1, MS2 | RT verified by standard | No |
| Identification level | Species level | Separation of isobaric/isomeric interferece confirmed | No |
| Polarity mode | Positive | Model for separation prediction | No |
| Type of positive (precursor)ion | [M+H]+ | Additional dimension/techniques | IMS |
| Fragments for identification | | CCS verified by standard | No |

| Fragment name |
|---|
| M-FA-TMA |

| | | | |
|---|---|---|---|
| Isotope correction at MS1 | No | How was/were the additional dimension(s) used? | For separation of isobaric/isomeric interferece in MS1 and MS2 dimensions |
| Isotope correction at MS2 | No | Was a model used to predict lipid molecule separation? | No |
| MS1 verified by standard | No | Lipid Identification Software | Skyline |
| MS2 verified by standard | No | Data manipulation | - |
| Background check at MS1 | No | Nomenclature for intact lipid molecule | No |
| Background check at MS2 | No | Nomenclature for fragment ions | N/A |
| Did you presume assumptions for identification? | No | Further identification remarks | - |
| Check on: | - | | |

#### 30) CAR, AC[M+H]+ / Lipid quantification

| | | | |
|---|---|---|---|
| Quantitative | No | Batch correction | No |
| Normalization to reference | No | Further quantification remarks | - |

#### 31) CAR, ANA[M+H]+ / Lipid identification

| | | | |
|---|---|---|---|
| Lipid class | CAR, ANA | RT verified by standard | No |
| MS Level for identification | MS1 | Separation of isobaric/isomeric interferece confirmed | No |
| Identification level | Species level | Model for separation prediction | No |
| Polarity mode | Positive | Additional dimension/techniques | IMS |
| Type of positive (precursor)ion | [M+H]+ | CCS verified by standard | No |
| Isotope correction at MS1 | No | How was/were the additional dimension(s) used? | For separation of isobaric/isomeric interferece |
| MS1 verified by standard | No | Was a model used to predict lipid molecule separation? | No |
| Background check at MS1 | No | Lipid Identification Software | Skyline |
| Did you presume assumptions for identification? | No | Data manipulation | - |
| Check on: | - | Nomenclature for intact lipid molecule | No |
| Limit of detection | No | Further identification remarks | - |

#### 31) CAR, ANA[M+H]+ / Lipid quantification

| | | | |
|---|---|---|---|
| Quantitative | No | Batch correction | No |
| Normalization to reference | No | Further quantification remarks | - |

#### 32) CAR, SE[M+NH4]+ / Lipid identification

| | | | |
|---|---|---|---|
| Lipid class | CAR, SE | Limit of detection | No |
| MS Level for identification | MS1, MS2 | RT verified by standard | No |
| Identification level | Species level | Separation of isobaric/isomeric interferece confirmed | No |
| Polarity mode | Positive | Model for separation prediction | No |
| Type of positive (precursor)ion | [M+NH4]+ | Additional dimension/techniques | IMS |
| Fragments for identification<br>**Fragment name**<br>-FA1(+HO)-Cholesterol(35) | | CCS verified by standard | No |
| Isotope correction at MS1 | No | How was/were the additional dimension(s) used? | For separation of isobaric/isomeric interferece in MS1 and MS2 dimensions |
| Isotope correction at MS2 | No | Was a model used to predict lipid molecule separation? | No |
| MS1 verified by standard | No | Lipid Identification Software | Skyline |
| MS2 verified by standard | No | Data manipulation | - |
| Background check at MS1 | No | Nomenclature for intact lipid molecule | No |
| Background check at MS2 | No | Nomenclature for fragment ions | N/A |
| Did you presume assumptions for identification? | No | Further identification remarks | - |
| Check on: | - | | |

#### 32) CAR, SE[M+NH4]+ / Lipid quantification

| | | | |
|---|---|---|---|
| Quantitative | No | Batch correction | No |
| Normalization to reference | No | Further quantification remarks | - |

#### 33) CAR, Cer[M+H]+ / Lipid identification

| | | | |
|---|---|---|---|
| Lipid class | CAR, Cer | Limit of detection | No |
| MS Level for identification | MS1, MS2 | RT verified by standard | No |
| Identification level | Molecular species level | Separation of isobaric/isomeric interferece confirmed | No |
| Polarity mode | Positive | Model for separation prediction | No |
| Type of positive (precursor)ion | [M+H]+ | Additional dimension/techniques | IMS |
| Fragments for identification | | CCS verified by standard | No |

| Fragment name |
|---|
| LCB(-CH3O2) |
| LCB(-H3O2) |
| LCB(-HO) |

| | | | |
|---|---|---|---|
| Isotope correction at MS1 | No | How was/were the additional dimension(s) used? | For separation of isobaric/isomeric interferece in MS1 and MS2 dimensions |
| Isotope correction at MS2 | No | Was a model used to predict lipid molecule separation? | No |
| MS1 verified by standard | No | Lipid Identification Software | Skyline |
| MS2 verified by standard | No | Data manipulation | - |
| Background check at MS1 | No | Nomenclature for intact lipid molecule | No |
| Background check at MS2 | No | Nomenclature for fragment ions | N/A |
| Did you presume assumptions for identification? | No | Further identification remarks | - |
| Check on: | - | | |

#### 33) CAR, Cer[M+H]+ / Lipid quantification

| | | | |
|---|---|---|---|
| Quantitative | No | Batch correction | No |
| Normalization to reference | No | Further quantification remarks | - |

#### 34) CAR, DG[M+NH4]+ / Lipid identification

| | | | |
|---|---|---|---|
| Lipid class | CAR, DG | Limit of detection | No |
| MS Level for identification | MS1, MS2 | RT verified by standard | No |
| Identification level | Molecular species level | Separation of isobaric/isomeric interferece confirmed | No |
| Polarity mode | Positive | Model for separation prediction | No |
| Type of positive (precursor)ion | [M+NH4]+ | Additional dimension/techniques | IMS |
| Fragments for identification | | CCS verified by standard | No |

| Fragment name |
|---|
| -FA1(-H)-(H2O+NH3) |
| -FA2(-H)-(H2O+NH3) |

| | | | |
|---|---|---|---|
| Isotope correction at MS1 | No | How was/were the additional dimension(s) used? | For separation of isobaric/isomeric interferece in MS1 and MS2 dimensions |
| Isotope correction at MS2 | No | Was a model used to predict lipid molecule separation? | No |
| MS1 verified by standard | No | Lipid Identification Software | Skyline |
| MS2 verified by standard | No | Data manipulation | - |
| Background check at MS1 | No | Nomenclature for intact lipid molecule | No |
| Background check at MS2 | No | Nomenclature for fragment ions | N/A |
| Did you presume assumptions for identification? | No | Further identification remarks | - |
| Check on: | - | | |

#### 34) CAR, DG[M+NH4]+ / Lipid quantification

| | | | |
|---|---|---|---|
| Quantitative | No | Batch correction | No |
| Normalization to reference | No | Further quantification remarks | - |

#### 35) CAR, LPE[M+H]+ / Lipid identification

| | | | |
|---|---|---|---|
| Lipid class | CAR, LPE | Limit of detection | No |
| MS Level for identification | MS1, MS2 | RT verified by standard | No |
| Identification level | sn Position | Separation of isobaric/isomeric interferece confirmed | No |
| Polarity mode | Positive | Model for separation prediction | No |
| Type of positive (precursor)ion | [M+H]+ | Additional dimension/techniques | IMS |
| Fragments for identification | | CCS verified by standard | No |
| Fragment name: -HG(PE,141) | | | |
| Isotope correction at MS1 | No | How was/were the additional dimension(s) used? | For separation of isobaric/isomeric interferece of MS1 and MS2 dimensions |
| Isotope correction at MS2 | No | Was a model used to predict lipid molecule separation? | No |
| MS1 verified by standard | No | Lipid Identification Software | Skyline |
| MS2 verified by standard | No | Data manipulation | - |
| Background check at MS1 | No | Nomenclature for intact lipid molecule | No |
| Background check at MS2 | No | Nomenclature for fragment ions | N/A |
| Did you presume assumptions for identification? | No | Further identification remarks | - |
| Check on: | - | | |

#### 35) CAR, LPE[M+H]+ / Lipid quantification

| | | | |
|---|---|---|---|
| Quantitative | No | Batch correction | No |
| Normalization to reference | No | Further quantification remarks | - |

#### 36) CAR, LPE[M+Na]+ / Lipid identification

| | | | |
|---|---|---|---|
| Lipid class | CAR, LPE | Limit of detection | No |
| MS Level for identification | MS1, MS2 | RT verified by standard | No |
| Identification level | sn Position | Separation of isobaric/isomeric interferece confirmed | No |
| Polarity mode | Positive | Model for separation prediction | No |
| Type of positive (precursor)ion | [M+Na]+ | Additional dimension/techniques | IMS |
| Fragments for identification | | CCS verified by standard | No |

| Fragment name |
|---|
| M-HG |
| M+ Na-az |

| | | | |
|---|---|---|---|
| Isotope correction at MS1 | No | How was/were the additional dimension(s) used? | For separation of isobaric/isomeric interferece in MS1 and MS2 dimensions |
| Isotope correction at MS2 | No | Was a model used to predict lipid molecule separation? | No |
| MS1 verified by standard | No | Lipid Identification Software | Skyline |
| MS2 verified by standard | No | Data manipulation | - |
| Background check at MS1 | No | Nomenclature for intact lipid molecule | No |
| Background check at MS2 | No | Nomenclature for fragment ions | N/A |
| Did you presume assumptions for identification? | No | Further identification remarks | - |
| Check on: | - | | |

#### 36) CAR, LPE[M+Na]+ / Lipid quantification

| | | | |
|---|---|---|---|
| Quantitative | No | Batch correction | No |
| Normalization to reference | No | Further quantification remarks | - |

#### 37) CAR, PC O[M+H]+ / Lipid identification

| | | | |
|---|---|---|---|
| Lipid class | CAR, PC O | Limit of detection | No |
| MS Level for identification | MS1, MS2 | RT verified by standard | No |
| Identification level | Molecular species level | Separation of isobaric/isomeric interferece confirmed | No |
| Polarity mode | Positive | Model for separation prediction | No |
| Type of positive (precursor)ion | [M+H]+ | Additional dimension/techniques | IMS |
| Fragments for identification | | CCS verified by standard | No |

| Fragment name |
|---|
| M-FA1 |
| M-oFA2 |

| | | | |
|---|---|---|---|
| Isotope correction at MS1 | No | How was/were the additional dimension(s) used? | For separation of isobaric/isomeric interferece in MS1 and MS2 dimensions |
| Isotope correction at MS2 | No | Was a model used to predict lipid molecule separation? | No |
| MS1 verified by standard | No | Lipid Identification Software | Skyline |
| MS2 verified by standard | No | Data manipulation | - |
| Background check at MS1 | No | Nomenclature for intact lipid molecule | No |
| Background check at MS2 | No | Nomenclature for fragment ions | N/A |
| Did you presume assumptions for identification? | No | Further identification remarks | - |
| Check on: | - | | |

#### 37) CAR, PC O[M+H]+ / Lipid quantification

| | | | |
|---|---|---|---|
| Quantitative | No | Batch correction | No |
| Normalization to reference | No | Further quantification remarks | - |

#### 38) CAR, PC O[M+Na]+ / Lipid identification

| | | | |
|---|---|---|---|
| Lipid class | CAR, PC O | Limit of detection | No |
| MS Level for identification | MS1, MS2 | RT verified by standard | No |
| Identification level | Molecular species level | Separation of isobaric/isomeric interferece confirmed | No |
| Polarity mode | Positive | Model for separation prediction | No |
| Type of positive (precursor)ion | [M+Na]+ | Additional dimension/techniques | IMS |
| Fragments for identification | | CCS verified by standard | No |

| Fragment name |
|---|
| M+Na-HG |
| M+Na-TMA |
| M-FA |

| | | | |
|---|---|---|---|
| Isotope correction at MS1 | No | How was/were the additional dimension(s) used? | For separation of isobaric/isomeric interferece in MS1 and MS2 dimensions |
| Isotope correction at MS2 | No | Was a model used to predict lipid molecule separation? | No |
| MS1 verified by standard | No | Lipid Identification Software | Skyline |
| MS2 verified by standard | No | Data manipulation | - |
| Background check at MS1 | No | Nomenclature for intact lipid molecule | No |
| Background check at MS2 | No | Nomenclature for fragment ions | N/A |
| Did you presume assumptions for identification? | No | Further identification remarks | - |
| Check on: | - | | |

#### 38) CAR, PC O[M+Na]+ / Lipid quantification

| | | | |
|---|---|---|---|
| Quantitative | No | Batch correction | No |
| Normalization to reference | No | Further quantification remarks | - |

#### 39) CAR, PE[M+H]+ / Lipid identification

| | | | |
|---|---|---|---|
| Lipid class | CAR, PE | Limit of detection | No |
| MS Level for identification | MS1, MS2 | RT verified by standard | No |
| Identification level | Molecular species level | Separation of isobaric/isomeric interferece confirmed | No |
| Polarity mode | Positive | Model for separation prediction | No |
| Type of positive (precursor)ion | [M+H]+ | Additional dimension/techniques | IMS |
| Fragments for identification | | CCS verified by standard | No |

| Fragment name |
|---|
| -HG(PE,141) |
| FA1 (+O) |
| FA2 (+O) |

| | | | |
|---|---|---|---|
| Isotope correction at MS1 | No | How was/were the additional dimension(s) used? | For separation of isobaric/isomeric interferece in MS1 and MS2 dimensions |
| Isotope correction at MS2 | No | Was a model used to predict lipid molecule separation? | No |
| MS1 verified by standard | No | Lipid Identification Software | Skyline |
| MS2 verified by standard | No | Data manipulation | - |
| Background check at MS1 | No | Nomenclature for intact lipid molecule | No |
| Background check at MS2 | No | Nomenclature for fragment ions | N/A |
| Did you presume assumptions for identification? | No | Further identification remarks | - |
| Check on: | - | | |

#### 39) CAR, PE[M+H]+ / Lipid quantification

| | | | |
|---|---|---|---|
| Quantitative | No | Batch correction | No |
| Normalization to reference | No | Further quantification remarks | - |

#### 40) CAR, PE[M+Na]+ / Lipid identification

| | | | |
|---|---|---|---|
| Lipid class | CAR, PE | Limit of detection | No |
| MS Level for identification | MS1, MS2 | RT verified by standard | No |
| Identification level | Molecular species level | Separation of isobaric/isomeric interferece confirmed | No |
| Polarity mode | Positive | Model for separation prediction | No |
| Type of positive (precursor)ion | [M+Na]+ | Additional dimension/techniques | IMS |
| Fragments for identification | | CCS verified by standard | No |

| Fragment name |
|---|
| M-HG |
| M+Na-HG |
| M+Na-az |
| M+Na-C2H5N-FA1 |
| M+Na-C2H5N-FA2 |

| | | | |
|---|---|---|---|
| Isotope correction at MS1 | No | How was/were the additional dimension(s) used? | For separation of isobaric/isomeric interference in MS1 and MS2 dimensions |
| Isotope correction at MS2 | No | Was a model used to predict lipid molecule separation? | No |
| MS1 verified by standard | No | Lipid Identification Software | Skyline |
| MS2 verified by standard | No | Data manipulation | - |
| Background check at MS1 | No | Nomenclature for intact lipid molecule | No |
| Background check at MS2 | No | Nomenclature for fragment ions | N/A |
| Did you presume assumptions for identification? | No | Further identification remarks | - |
| Check on: | - | | |

#### 40) CAR, PE[M+Na]+ / Lipid quantification

| | | | |
|---|---|---|---|
| Quantitative | No | Batch correction | No |
| Normalization to reference | No | Further quantification remarks | - |

#### 41) CAR, SM[M+H]+ / Lipid identification

| | | | |
|---|---|---|---|
| Lipid class | CAR, SM | Limit of detection | No |
| MS Level for identification | MS1, MS2 | RT verified by standard | No |
| Identification level | Molecular species level | Separation of isobaric/isomeric interferece confirmed | No |
| Polarity mode | Positive | Model for separation prediction | No |
| Type of positive (precursor)ion | [M+H]+ | Additional dimension/techniques | IMS |
| Fragments for identification | | CCS verified by standard | No |

| Fragment name |
|---|
| LCB(-H3O2) |

| | | | |
|---|---|---|---|
| Isotope correction at MS1 | No | How was/were the additional dimension(s) used? | For separation of isobaric/isomeric interferece in MS1 and MS2 dimensions |
| Isotope correction at MS2 | No | Was a model used to predict lipid molecule separation? | No |
| MS1 verified by standard | No | Lipid Identification Software | Skyline |
| MS2 verified by standard | No | Data manipulation | - |
| Background check at MS1 | No | Nomenclature for intact lipid molecule | No |
| Background check at MS2 | No | Nomenclature for fragment ions | N/A |
| Did you presume assumptions for identification? | No | Further identification remarks | - |
| Check on: | - | | |

#### 41) CAR, SM[M+H]+ / Lipid quantification

| | | | |
|---|---|---|---|
| Quantitative | No | Batch correction | No |
| Normalization to reference | No | Further quantification remarks | - |

#### 42) CAR, TG[M+NH4]+ / Lipid identification

| | | | |
|---|---|---|---|
| Lipid class | CAR, TG | Limit of detection | No |
| MS Level for identification | MS1, MS2 | RT verified by standard | No |
| Identification level | Molecular species level | Separation of isobaric/isomeric interferece confirmed | No |
| Polarity mode | Positive | Model for separation prediction | No |
| Type of positive (precursor)ion | [M+NH4]+ | Additional dimension/techniques | IMS |
| Fragments for identification | | CCS verified by standard | No |

| Fragment name |
|---|
| -FA3(+HO)-(NH3) |
| -FA2(+HO)-(NH3) |
| -FA1(+HO)-(NH3) |
| FA3 |
| FA2 |
| FA1 |

| | | | |
|---|---|---|---|
| Isotope correction at MS1 | No | How was/were the additional dimension(s) used? | For separation of isobaric/isomeric interferece of MS1 and MS2 dimensions |
| Isotope correction at MS2 | No | Was a model used to predict lipid molecule separation? | No |
| MS1 verified by standard | No | Lipid Identification Software | Skyline |
| MS2 verified by standard | No | Data manipulation | - |
| Background check at MS1 | No | Nomenclature for intact lipid molecule | No |
| Background check at MS2 | No | Nomenclature for fragment ions | N/A |
| Did you presume assumptions for identification? | No | Further identification remarks | - |
| Check on: | - | | |

#### 42) CAR, TG[M+NH4]+ / Lipid quantification

| | | | |
|---|---|---|---|
| Quantitative | No | Batch correction | No |
| Normalization to reference | No | Further quantification remarks | - |

#### 43) CAR, CL[M+H]+ / Lipid identification

| | | | |
|---|---|---|---|
| Lipid class | CAR, CL | RT verified by standard | No |
| MS Level for identification | MS1 | Separation of isobaric/isomeric interferece confirmed | No |
| Identification level | Species level | Model for separation prediction | No |
| Polarity mode | Positive | Additional dimension/techniques | IMS |
| Type of positive (precursor)ion | [M+H]+ | CCS verified by standard | No |
| Isotope correction at MS1 | No | How was/were the additional dimension(s) used? | For separation of isobaric/isomeric interferece |
| MS1 verified by standard | No | Was a model used to predict lipid molecule separation? | No |
| Background check at MS1 | No | Lipid Identification Software | Skyline |
| Did you presume assumptions for identification? | No | Data manipulation | - |
| Check on: | - | Nomenclature for intact lipid molecule | No |
| Limit of detection | No | Further identification remarks | - |

#### 43) CAR, CL[M+H]+ / Lipid quantification

| | | | |
|---|---|---|---|
| Quantitative | No | Batch correction | No |
| Normalization to reference | No | Further quantification remarks | - |
